# Mutant analysis of Kcng4b reveals how the different functional states of the voltage-gated potassium channel regulate ear development

**DOI:** 10.1101/2023.06.26.546501

**Authors:** Justyna Jędrychowska, Vitya Vardanyan, Milosz Wieczor, Antoni Marciniak, Jacek Czub, Razieh Amini, Ruchi Jain, Hongyuan Shen, Hyungwon Choi, Jacek Kuznicki, Vladimir Korzh

## Abstract

The voltage gated (Kv) slow-inactivating delayed rectifier channel regulates the development of hollow organs of the zebrafish. The functional tetramer consists of an electrically active subunit (Kcnb1, Kv2.1) and a modulatory silent subunit (Kcng4b, Kv6.4). The two mutations in zebrafish *kcng4b - kcng4b-C1* and *kcng4b-C2* (Gasanov et al., 2021) - have been studied during ear development using electrophysiology, developmental biology and *in silico* structural modelling. *kcng4b-C1* mutation causes a C-terminal truncation characterized by mild Kcng4b loss-of-function (LOF) manifested by failure of kinocilia to extend and formation of ectopic otoliths. In contrast, the *kcng4b-C2^-/-^* mutation causes the C-terminal domain to elongate and the ectopic seventh transmembrane (TM) domain to form, converting the intracellular C-terminus to an extracellular one. Kcng4b-C2 acts as a Kcng4b gain-of-function (GOF) allele. Otoliths fail to develop and kinocilia are reduced in *kcng4b-C2^-/-^*. These results show that different mutations of the silent subunit Kcng4 can affect the activity of the Kv channel and cause a wide range of developmental defects.

## Introduction

Development is regulated by biochemical gradients, transcriptional networks, and bioelectrical signals generated by ion channels and pumps (Levin, 2014). Voltage-gated potassium (K_v_) channels play an important role in the maintenance of the membrane potential. Their activity is tightly regulated. This is critical for the development and function of various cell lineages (Jędrychowska and Korzh, 2019; Shen et al., 2016; Vacher et al., 2008). In humans, mutations in K_v_ channels-encoding genes cause various of diseases (Felix, 2000; Simons et al., 2015). For example, Kv2.1 (KCNB1) dysfunction has been associated with epilepsy (Bar et al., 2020; de Kovel et al., 2017; Kang et al., 2019; Latypova et al., 2017; Marini et al., 2017; Torkamani et al., 2014). This disease is likely caused by developmental abnormalities (Hawkins et al., 2021; Jędrychowska and Korzh, 2019; Speca et al., 2014).

Kv2 channels consists of the electrically active (KCNB1) and modulatory (or silent, KCNG4) α-subunits. Subunit monomers assemble functional channel heterotetramers. The stoichiometry of the subunits in any given tetramer is variable, which adds to the functional complexity of K_v_ channel (Bocksteins, 2016; Möller et al., 2020). The genes encoding the subunits are expressed in the cell lineage and tissue specific pattern. Antagonizing activities of different subunits of K_v_ channels establish a regulated balance *in vitro* (Bocksteins and Snyders, 2012; Ottschytsch et al., 2005) and *in vivo* (Jedrychowska et al., 2021; Jędrychowska and Korzh, 2019; Shen et al., 2016).

Most human KCNB1 mutants are heterozygotes carrying a single amino acid substitution. They develop early-induced epileptic encephalopathy (EIEE26), which is characterized by severe cognitive impairment (de Kovel et al., 2017; Jędrychowska and Korzh, 2019; Niday and Tzingounis, 2018; Saitsu et al., 2015; Srivastava et al., 2014; Thiffault et al., 2015; Torkamani et al., 2014). In comparison, mammalian Kcng4 analyses are much more limited. The GWAS study suggests that defects of KCNG4 may be associated with migraine, multiple sclerosis and modulation of the uterine nociceptors excitability during labor (Lafrenière and Rouleau, 2012; Lee et al., 2020; Vilariño-Güell et al., 2019), while targeted deletion of Kcng4 in mice results in male sterility (Regnier et al., 2017) and hydrocephalus (https://www.mousephenotype.org/data/genes/MGI:1913983#phenotypesTab).

Some KCNB1 heterozygotes have developmental delay and speech defects (Bar et al., 2020; Latypova et al., 2017; Marini et al., 2017), which may be due to KCNB1 gain-of-function (GOF) (Veale et al., 2022). Speech develops with the contribution of many inputs, including the hearing of sound (Anne et al., 2017). The orientation of aquatic and terrestrial animals in three-dimensional (3D) space depends on the inner ear, which coordinates the hearing of sound and the sense of balance. The latter relies on the vestibular system - three semicircular canals, where mechanosensory cells detect changes of fluid dynamics depending on the body position in 3D space and gravity. Sensation depends on the interaction of sensory cells with aggregates of biominerals such as otoconia in mammals or otoliths (utricle, saccule, and lagena) in non-mammalian vertebrates. Otoliths are composed of calcium carbonate and specialized proteins [Otopetrin (Otop1), Starmaker-like 1 (Sml-1), etc.] (Bezares-Calderón et al., 2020; Go et al., 2010; Hudspeth and Logothetis, 2000; Liedtke et al., 2000; Rózycka et al., 2014). Otolith formation depends on motile cilia that move the K^+^-enriched endolymph and tether cilia to which the otoliths attach (Colantonio et al., 2009; Riley et al., 1997; Yu et al., 2011; Wu et al, 2011).

Genetic analysis of ear development in various vertebrates, such as zebrafish, medaka, and mice, has identified a few genes that regulate ear and otolith development. Deficiency of these genes causes reduction, fusion, displacement, or absence of otoliths, resulting in hearing and balance defects (Chatterjee and Lufkin, 2011; Dror and Avraham, 2009; Malicki et al., 1996; Söllner et al., 2004; Steel and Brown, 1994; Whitfield et al., 1996). For example, the proton-selective ion channel Otopetrin1 (otop1) is required for the formation of otoconia/otoliths (Hughes et al., 2004; Söllner et al., 2004; Whitfield, 2020). The ciliary cytoskeleton and intraflagellar trafficking are essential for cilia extension (Choksi et al., 2014; Pathak and Drummond, 2009). Failure of cilia extension has been associated with language defects (Stawicki et al., 2016).

We have shown that zebrafish lacking the electrically-active subunit of Kcnb1 develop microcephaly, reduced otocyst and otoliths, and deformed kinocilia (Jedrychowska et al., 2021; Shen et al., 2016). In contrast, zebrafish *Kcng4b* mutants develop hydrocephalus (Shen et al., 2016).

*kcng4b* is expressed only during development in the brain ventricular system (BVS), ear, *etc.* (Shen et al., 2016). The developmental role of Kcng4 in the ear is still not fully understood. To address this issue, we generated two C-terminal *kcng4b* mutants (Gasanov et al., 2021). Unexpectedly, these cause opposite developmental defects ear development, likely because they represent the gain-of-function and loss-of -function mutant versions of Kcng4.

## Materials and Methods

### Animals

The wild-type (AB) and mutant zebrafish (*Danio rerio*) lines were maintained in the Zebrafish Facility of the International Institute of Molecular and Cell Biology in Warsaw (license No. PL14656251) according to the standard procedures and ethical practices (Westerfield, 2007). All experiments with zebrafish embryos/larvae were performed in accordance with the rules of the Polish Laboratory Animal Science Association and European Communities Council Directive (63/2010/EEC). Developmental stages (in hours, hpf, and days post fertilization, dpf) were described according to (Kimmel et al., 1995). The transgenic line, *Tg(Brn3c:GAP43-GFP)^s356t^* (Xiao et al., 2005), which expresses cytosolic EGFP in inner ear and lateral line mechanosensory hair cells, was used in this study. The generation of two *kcng4b* mutations, C1 and C2, by CRISPR-Cas9 has been described previously (Gasanov et al., 2021).

*Xenopus laevis* frogs (Nasco, Chicago, IL) were maintained in the animal facility of the Institute of Molecular Biology NAS RA (permission number 13-1F322) according to the guidelines of the local animal welfare authorities. All the quantified data are presented as mean ± SD.

### Heterologous expression of K_v_ channels in *X. laevis* oocytes and electrophysiological measurements

Human K_v_2.1 wild-type channel was co-expressed with zebrafish Kcng4b wild-type protein its mutant forms in the *X. laevis* oocytes. *Kcng4b-wt*, *Kcng4b-C1*, *Kcng4b-C2*, and *Kcng4b-trunc* constructs were subcloned from pTnT vector into the oocyte pGEM-He-juell oocyte vector using *Bam*HI and *Hind*III sites, between the *X. laevis* β-globin 5’-and 3’-untranslated sequences to increase the efficiency of expression. The *in vitro* transcription was made using the mMessage mMachine Kit (Ambion, USA). Quality of synthesized RNA was checked in agarose gel-electrophoresis. The synthesized mRNAs were quantified with RNA 6000 Nano kit (Agilent Technologies, USA).

*X. laevis* oocytes were surgically removed and defolliculated using 2 mg/ml collagenase (Sigma-Aldrich, USA) in Ca^2+^-free OR_2_ solution: 82.5 mM NaCl, 2 mM KCl, 1 mM MgCl_2_, 5 mM HEPES, pH 7.5. Stage V and VI oocytes were selected for mRNA (12 pg/oocyte) injection using manual nano-injector (Drummond Scientific, USA). Injected oocytes were incubated in OR_2_ solution supplemented with 2 mM CaCl_2_, 5 mM sodium-pyruvate and 50 µg/ml gentamycin for 1-7 days prior to electrophysiological recording. The molecular ratio of K_v_2.1/Kcng4b subunits for co-expression experiments was 1:1.

Oocytes expressing K_v_2.1 and K_v_2.1/Kcng4 channels were analyzed by two-electrode voltage clamp (TEVC) method at room temperature. TURBO TEC-03X (NPI Electronic GmbH, Germany) amplifiers linked to Pulse software (HEKA Electronics, Germany) through ITC-16 digitizer were used for data acquisition and monitoring. Pipettes were pulled from borosilicate glass (WPI, Germany) using the PB/7 puller (Narishige, Japan). Tips of the recording electrodes were prefilled with 1% agar-3M KCl and backfilled with 3M KCl solutions to prevent KCl leakage into the cells. Recordings were performed in modified ND96 solution containing: 75 mM NaCl, 5 mM KCl, 2 mM CaCl_2_, 1 mM MgCl_2_, 5 mM HEPES, pH 7.5. Membrane voltage was continuously monitored during the recordings.

Conductance-voltage relations for wild-type and mutant K_v_ channels were determined from tail currents. The Pulsfit (HEKA), Igor Pro 6.0 (WaveMetrics, Lake Oswego, OR, USA) and Kaleidagraph 4.0 (Synergy Software, Reading, PA, USA) software for data analysis were used. Non-specific leak current was estimated by averaging the currents of 4-7 water-injected oocytes in parallel to each experiment. The average current was subtracted for GV and steady-state inactivation analysis. Normalized data points were fitted with Boltzmann function of the form G/G_max_=1/(1+e(^(V1/2-V)/K)*F/RT^) for GVs and I_test_/I_cont_=((1-offset)/(1+e(^(V1/2-V)/K)*F/RT^)) +offset for steady-state inactivation, V_1/2_ is the midpoint voltage of activation and inactivation, respectively, K is the slope constant, F, R and T have their usual meanings.

### **C**omputer modelling of Kcng4b/**K_v_6.4b** mutant proteins

The isolated S7 helix model was generated by Tasser web server (Yang et al., 2014) using a sequence of 34 amino acids predicted to form a TM helix. The structure of the K_v_1.2-2.1 paddle chimera channel (PDB id: 2R9R) was used as a template for the tetramer model after discarding beta-2 chains. Two opposing monomers served as the basis for modelling Kcng4b, whereas the other two were left unchanged. When constructing the Kcng4b monomer structures, the S7 helix was added based on the isolated helix model with constraints on the distances between the ends of S7 and S1 helices. The Modeller software (Webb and Sali, 2016) was used for these operations. Both models were embedded in membranes using MembraneBuilder from the CHARMM-GUI web server (Wu et al., 2014) and simulated using Gromacs 5.1.4 software (Abraham et al., 2015) with the CHARMM36m force field at physiological ionic strength (0.15 M KCl). Each system was minimized using the standard MembraneBuilder protocol and simulations were performed under isothermal-isobaric (NPT) conditions with semi-isotropic pressure coupling, temperature maintained at 310 K, and using the standard particle mesh Ewald (PME) cutoff of 1.2 nm. The single helix was simulated for 1.3 μs in the liquid-ordered (Lo) phase – 50 dipalmitoylphosphatidylcholine (DPPC) and 20 cholesterol molecules per leaflet – and 1.5 μs in liquid-disordered (Ld) phase – 50 phosphatidylcholine (POPC) molecules per leaflet. The tetramer was first simulated with positional restraint on Ca for 250 ns to ensure its stability, then with root-mean-square deviation (RMSD) restraint: RMSD of every 3^rd^ Ca of helical parts of the TM domain was restrained at 0.15, in reference to the Modeller model. The restraint force constant was kept at 100000 for 200 ns, 25000 for 400 ns and 5000 for 450 ns. Finally, the model was simulated without restraints for 1.2 μs. We used Gromacs for position restraints and plumed for RMSD restraints (Tribello et al 2014).

Mean square displacement (MSD) and angle calculations were performed using the MDTraj library for Python (McGibbon et al., 2015). Only backbone atoms were used for both analyses.

### Microscopy

#### Live imaging

Zebrafish embryos were raised in E3 medium (2.5 mM NaCl, 0.1 mM KCl, 0.16 mM CaCl_2_, and 0.43 mM MgCl_2_) with the addition of 0.2 mM 1-phenyl-2-thiourea (PTU, Merck, Germany) to block pigmentation. At selected developmental stages, embryos were manually dechorionated, anesthetized with 0.02% tricaine (Sigma-Aldrich, USA) and oriented by embedding in 2 % methylcellulose (Merck, Germany) on the glass slides. A research stereomicroscope SMZ25 (Nikon, Japan) was used to image acquisition.

Vital dye staining of lateral line hair cells of the 4 dpf embryos was performed using DASPEI (2-(4-(dimethylamino) styryl)-N-ethylpyridinium iodide) (Sigma-Aldrich, USA, D0815), 0.8 µg/ml, and Calcium Crimson, AM ester (Thermo Fisher, USA, C3018), 6,8 µg/ml in E3 medium, 28° C, 15 min. 3 μm FM 1-43FX was injected into an ear of 5 dpf larvae as described in (Smith et al., 2020).

#### Light-sheet and confocal fluorescence microscopy imaging

*In vivo* imaging and imaging of fixed specimens were performed as previously described (García-Lecea et al., 2017; Jedrychowska et al., 2021). For imaging of fixed material, 0.8 % low-melting agarose in phosphate buffer saline (PBS) was used instead of E3 0.02 % tricaine medium. Zeiss Lightsheet Z. 1 with W Plan-Apochromat 20x/1.0 UV-VIS (for *in vivo* imaging), 40x/1.0 UV-VIS or 63x/1.0 UV-VIS (for fixed embryos) objectives were used, transmitted LED light was also used to obtain the high-resolution bright-field images of zebrafish ear. Confocal imaging was performed using the Zeiss LSM 800 inverted microscope with Airyscan (Carl Zeiss, Germany). 405, 493 and 575 nm lasers were used to excite fluorescence with emission detected using emission filters (425–430, 514–530 and 592–625 nm BP), respectively. Data were saved in the LSM or CZI format and processed using ZEN (Zeiss) or ImageJ 1.51n (Fiji) software. Maximum intensity or sum slices projections were generated for each z-stack. Brightness / contrast adjustments and resizing were performed using FastStone viewer 7.4 (FastStone Soft).

### Immunohistochemistry and *in situ* hybridization

Whole-mount immunohistochemistry was performed according to a previously established protocol (Korzh et al., 1998). A 1:100 dilution of mouse monoclonal anti-acetylated tubulin antibody (Sigma-Aldrich, USA, T7451) was used as the primary antibody and 1:2000 dilution of donkey anti-mouse Alexa Fluor 488 (Thermo Fisher, USA, R37114) was used as the secondary antibody. Alexa Fluor 568 Phalloidin (Thermo Fisher, USA, A12380) staining (1:200) was performed simultaneously with the secondary antibody incubation (Tanimoto et al., 2011). DAPI (4’,6’-Diamidine-2’-phenylindole dihydrochloride, Abcam, USA) was added to some samples along with secondary antibodies to final concentration 10 µM. RNA probes for *in situ* hybridization were transcribed *in vitro* from the pTnT vector (Promega, Madison, WI, USA) with sequences of *kcnb1*, *kcng4b*, *otop1*, *slc12a2* and *atp2b1a* cloned in 3’-5’direction cDNA using Digoxigenin-labelled uridine triphosphate (DIG-UTP). The probe against *kcnb1* was synthesized from a previously described plasmid (Shen et al., 2016). The synthesis of the *otop1* and *atp2b1a* probes is described previously (Jedrychowska et al., 2021), while the *kcng4b* and *slc12a2* probes were generated from pKcng4b-probe/pTnT and slc12a2-probe/pTnT vectors. These vectors were prepared by PCR using the following primers: Kcng4b-p-fvd (5’- ATATAT<u>GAATTC</u>ATGCCCATCATCAGCAATGCTAACC) and Kcng4b-p-rev (5’- ATATAT<u>GAATTCC</u>TGAAGCCCTTGTCCTGTGCAAGAGG), Slc12a2-fvd (5’- ATATAT<u>GAATTC</u>ATGTCGGCGTCACCTCCAATCTCCG) and Slc12a2-rev (5’- ATATAT<u>GAATTCC</u>TGATCCATCCGAACTTCACCACTCC), as well as zebrafish cDNA as a template. The PCR product was cloned into a pTnT vector using the *Eco*RI restriction site (underlined in the primer sequences) in both orientations to get the sense and antisense probes.

### Transcriptome analysis by RNAseq and bioinformatics

150 embryos for each sample were collected at 24 hpf [wild type control and Kcng4b-deficient crispants and morphants (Shen et al., 2016)] and RNA preparation was performed as previously published (Shen et al., 2017; Winata et al., 2013). mRNA levels were quantified using a standard pipeline based on TopHat-cufflinks59 with the *Danio rerio* gene annotation file (assembly GRCz10) from the Ensembl database. For differential expression analysis, gene-level FPKM values were converted into log ratios (KO/WT, base 2) and the threshold for differential expression to meet 1% FDR was determined using a mixture model-based approach called EBprot60. The transcripts whose largest FPKM values were below 25 percentiles of all FPKM values across the sample were removed from further analysis as unreliable. For genes whose expression changed in crispants and morphants compared to WT, a gene was considered differentially expressed if the quantified FPKM value (Trapnell et al., 2010) was above the 25 percentiles of the whole transcriptome in the given sample. Functional enrichment test (Gene Ontology) was performed using an in-house program that calculates the significance of enrichment by hypergeometric test, using the GO annotation of genes from ZFIN database (http://zfin.org). The RNAseq data have been deposited online (GEO submission no. GSE194272).

### qRT-PCR

The SsoAdvanced Universal SYBR Green Supermix and CFX Connect real-time PCR system (Bio-Rad, USA) was used for the Quantitative Real-Time Polymerase Chain Reaction (qRT-PCR) according to the manufacturer’s instructions. The TRIzol-chloroform RNA extraction protocol (Sigma-Aldrich, USA) was used to extract total RNA from 20-50 zebrafish embryos at 28, 48, and 72 hpf. RNA concentration was assessed using a NanoDrop™ 2000 spectrophotometer (Thermo Scientific, USA), and cDNA was synthesized from 1 µg of RNA using the iScript™ Reverse Transcription kit (Bio-Rad, USA). qRT-PCR was performed using gene-specific primers for *eef1a1l1*, the reference reaction for the housekeeping gene, and other primers (Suppl. Table 1) were used to amplify mRNAs of interest selected based on results of the RNAseq analysis described above.

The threshold cycle of each target and reference gene amplification in control and mutant embryos was determined automatically by LightCycler® 96 Software (Roche Diagnostics, USA). Fold change in mutants versus controls was calculated with delta-delta-C(t) method and Student’s two tailed t-test with respect to the mismatch control was used to determine statistical significance. Statistical analysis including standard deviation calculation, was performed using Microsoft Excel (Microsoft, USA) and GraphPad Prism 5 (GraphPad, USA) software. Melting curve analysis and agarose gel electrophoresis were performed as product specificity controls, while the reaction efficiency (E) was calculated separately for each gene analyzed. All RNA samples used as the templates were extracted independently from three or four different groups of embryos, and each was analyzed by qRT-PCR at least twice after cDNA synthesis. The value above 1.0 corresponds to an increase of mRNA level, the value below 1.0 corresponds to a decrease of mRNA level and “n.c.” - “not changed” is the same value as in the controls.

## Results

*kcng4b* plays the developmental role in the brain ventricular system (BVS) (Shen et al., 2016). Here we focused on the analysis of Kcng4b function during ear development, where the third (ectopic) otolith was previously detected in the insertional mutant *kcng4b-trunc* (*kcng4b^sq301^*) (Suppl. Fig. 1). In absence of this mutant line, which has been lost, to study the developmental role of Kcng4b, we generated two novel Kcng4b mutant alleles (*kcng4b-C1, kcng4b-C2*) (Fig. 1) both of which were lethal as homozygotes.

**Fig. 1.**
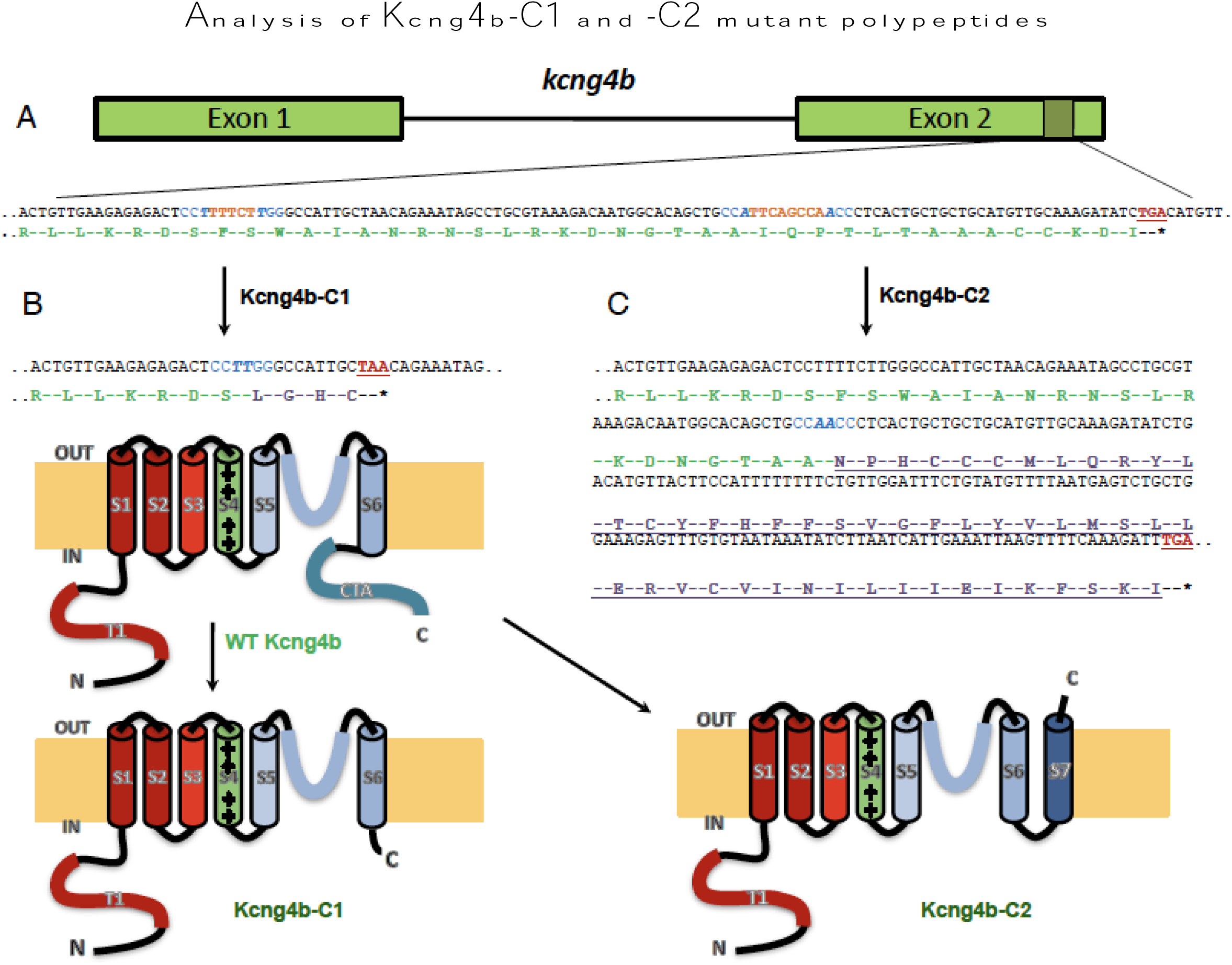
*kcng4b-C1* and *kcng4b-C2* mutations differently affect Kcng4b’s C-terminal. A - *kcng4b* nucleotide sequence is in black, stop-codons are in red, gRNAs’ PAM-sequences are in blue, and deleted nucleotides are in orange. The Kcng4b amino-acid sequence is in green, the modified one in lilac. B - mutation *kcng4b-C1* caused Kcng4b C-terminal truncation. C - mutation *kcng4b-C2* caused an extension of Kcng4b polypeptide and formation of an additional C-terminal TM domain (S7, underlined). T1 and CTA – N- and C-terminal cytoplasmic domains, S1-S7 – TM domains. IN, OUT – intracellular and extracellular space.

## Computational analysis of Kcng4b mutants

The deletion of five nucleotides in the second exon of *kcng4b* in the *kcng4b-C1* mutant caused the open reading frame (ORF) shift (Fig. 1A, B). The C-terminal of Kcng4b polypeptide was prematurely terminated with the loss of 29 C-terminal amino-acid residues (Gasanov et al., 2021) (Fig. 1B). In the second exon of *kcng4b-C2* mutant, eight nucleotides were deleted, resulting in the ORF shift that replaced 14 C-terminal amino acids with 49 ectopic ones (Gasanov et al., 2021)(Fig. 1C). To verify the structure of mutant polypeptides, we analyzed the putative amino acid sequences *in silico* using TMHMM software (https://services.healthtech.dtu.dk/service.php?TMHMM-2.0) and molecular modelling. These predicted no significant changes in the structure of the *kcng4b-C1* mutant other than its C-terminal truncation (Fig. 2A, B; 3A). In contrast, in the *kcng4b-C2* mutant, the ectopic seventh TM domain (S7) was predicted in the C-terminal of the putative Kcng4b-C2 polypeptide (Fig. 2C; 3B). Such a domain may flip the cytoplasmic C-terminal to an extracellular one, which will have a significant impact on the heterotetramer structure and function.

**Fig. 2.**
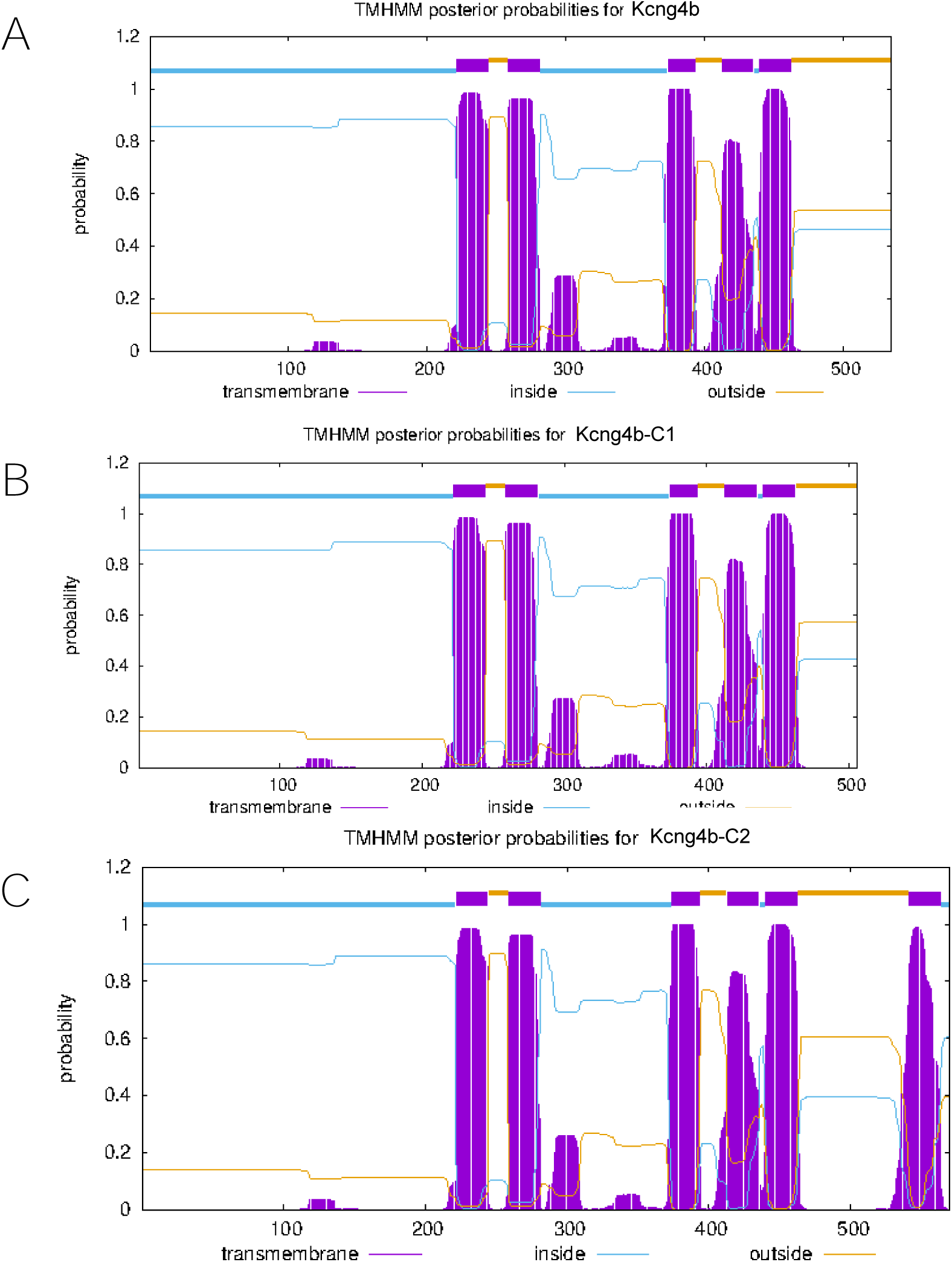
Hydrophobicity profiles illustrated an appearance in the Kcng4b-C2 of additional TM domain. A - Kcng4b (wild type), B - Kcng4b-C1; C - Kcng4b-C2.

To explore this possibility molecular dynamics simulations of the S7 helix in model lipid membranes were performed (see M&M for details). Since the behavior of TM helices is phase-dependent, we used POPC lipids as a model for liquid-disordered (Ld) domains, and DPPC lipids with 40% cholesterol as a model of liquid-ordered (Lo) domains (Fig. 3C). The S7 behaved differently in these cases: in the thinner Ld phase, the helix tilted by 44 degrees to better accommodate its hydrophobic thickness in the membrane hydrocarbon core, while in the much thicker Lo phase, it remained stably upright with average tilt of 11 degrees (Fig. 3C).

**Fig. 3.**
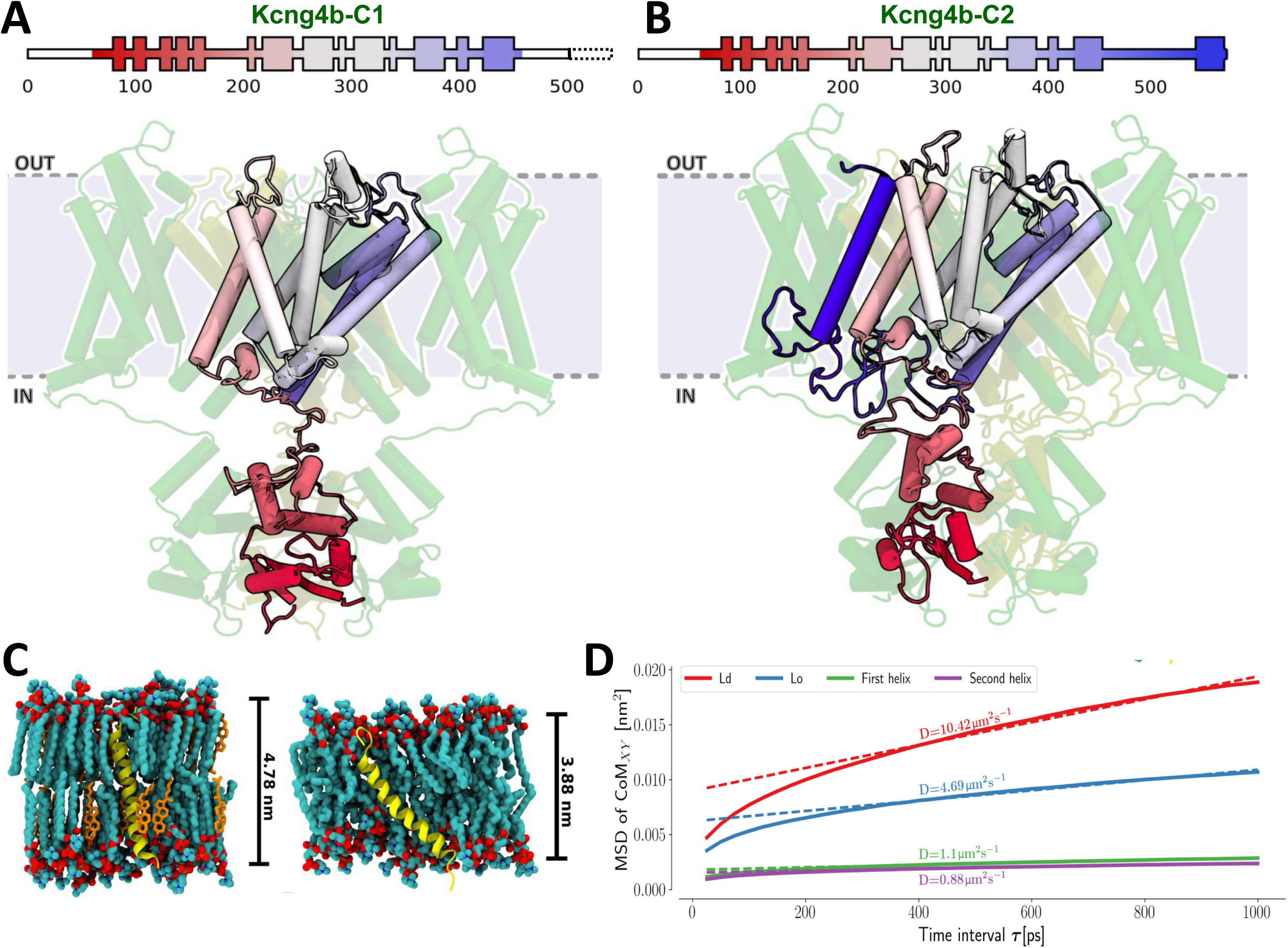
Modeling of Kcng4b/Kv6.4 mutant monomers, Kcng4b-C1 (A) and Kcng4b-C2 (B). IN, OUT – internal and external space. The deleted C-terminal region of Kcng4b-C1 is indicated by the dashed box. The additional TM domain of Kcng4b-C2 (S7) is in dark blue. C - Representations of isolated helix systems of Kcng4b-C2; S7 in a thicker, more stiff liquid ordered (Lo) phase (left), S7 in a thinner, looser liquid disordered (Ld) phase (right). D - Plots of mean square displacement (MSD) against time interval τ. Helices described as first and second refer to the S7s from two different Kcng4b/Kv6.4 monomers in the tetramer. Here, MSD measures displacement of the center of mass of the helix in membrane plane within the time interval τ. The slope of the MSD at long times is related to the diffusion coefficient D, equivalent to one obtained by standard experimental methods, e.g., single-molecule tracking.

To explore whether the S7 helix forms a stable complex within the K_v_2.1 (Kcnb1)-K_v_6.4 (Kcng4) heterotetramer, we set out to simulate the entire TM portion of the channel. Since a structure of an entire K_v_6.4 has not been solved, we used the K_v_1.2-K_v_2.1 paddle chimera channel (PDB ID: 2R9R) as a basis for the homology model in approach used previously (Matthies et al., 2018).

To show that the S7 helix remains stably bound to the channel, we compared the diffusion coefficients of the S7 helix either in pure lipid bilayers or in the context of the heterotetramer model by plotting mean squared displacement (MSD) against the time interval τ (Fig. 3D). The helices in the heterotetramer model are much less mobile than in the pure bilayer models, indicating that the S7 helix remains bound to the rest of the TM domains. Although we were unable to predict its exact rotational orientation with respect to the tetramer, the fact that its protein-protein interface has not been evolutionarily optimized strongly suggests that there is no single high-affinity binding site, and that the location of the additional helix is largely determined by an entropic tendency to displace lipids from the lateral surface of the protein where they undergo conformational ordering (Fig. 3B).

While the 35 amino acids of Kcng4b-C2 mutant discussed above clearly form a helix, this region is preceded by the 14 amino acids sequence whose secondary structure remains unknown. However, since this sequence results from the same ORF shift, in the absence of strong predictions, we found it reasonable to assume that this region is unstructured. Indeed, in our simulations, this region remained unstructured for a total of 2.25 µs, forming tangled loops with no clearly defined secondary structure (Fig. 3B).

Overall, both models confirm that S7 has the properties of a TM helix. The computer modelling revealed a possibility of formation of the Kcnb1-Kcng4 heterotetrameric potassium voltage-gated channel with the participation of Kcng4b-C2 and without obvious effect on channel structure (Fig. 3B; Suppl. Fig. 2) other that the flip of the C-terminal of Kcng4b-C2, making it an extracellular one. The *kcng4b-C2* mutation likely affects an interaction of the Kcng4b (Kv6.4) C-terminal and Kcnb1 (Kv2.1) N-terminal (Bocksteins et al., 2014).

### Kcng4b mutations differentially affect electrophysiological properties of Kv2.1 channels

To assess the functional effect of the *kcng4b-C1* and *kcng4b-C2* mutations on the expression and biophysical properties of Kv2.1 channel, we co-expressed the *kcng4b* mutant variants and wild-type mRNA (*kcng4b-wt*) with human *KCNB1* mRNA in *Xenopus* oocytes and compared the current densities and biophysical properties of the resultant channels. Co-expression of *kcng4b-wt* with *KCNB1* resulted in a reduction of the Kv2.1 current density and shifted the steady-state inactivation curve toward the hyperpolarized potentials (Fig. 4, Table 1). These findings were consistent with the previous results obtained by co-expression of *KCNG4* with *KCNB1* in *Xenopus* oocytes (Möller et al., 2020) as well as in mammalian cells (Shen et al., 2016). Co-injection of *KCNB1* and *kcng4b-C1* mRNA had a very similar effect on the biophysical properties of Kv2.1 as co-expression with *kcng4b-wt* (Fig. 4, Table 1). However, compared to the modulatory effect of Kcng4b-wt on Kv2.1 activity, Kcng4b-C1 is slightly but statistically significantly less efficient (Table 1) as indicated by the analysis of peak currents at +60 mV. The almost identical biophysical properties of Kv2.1/Kcng4b-C1 compared to Kv2.1/Kcng4b-wt suggest that Kcng4b-C1 forms typical heterotetramers with Kv2.1.

**Fig. 4.**
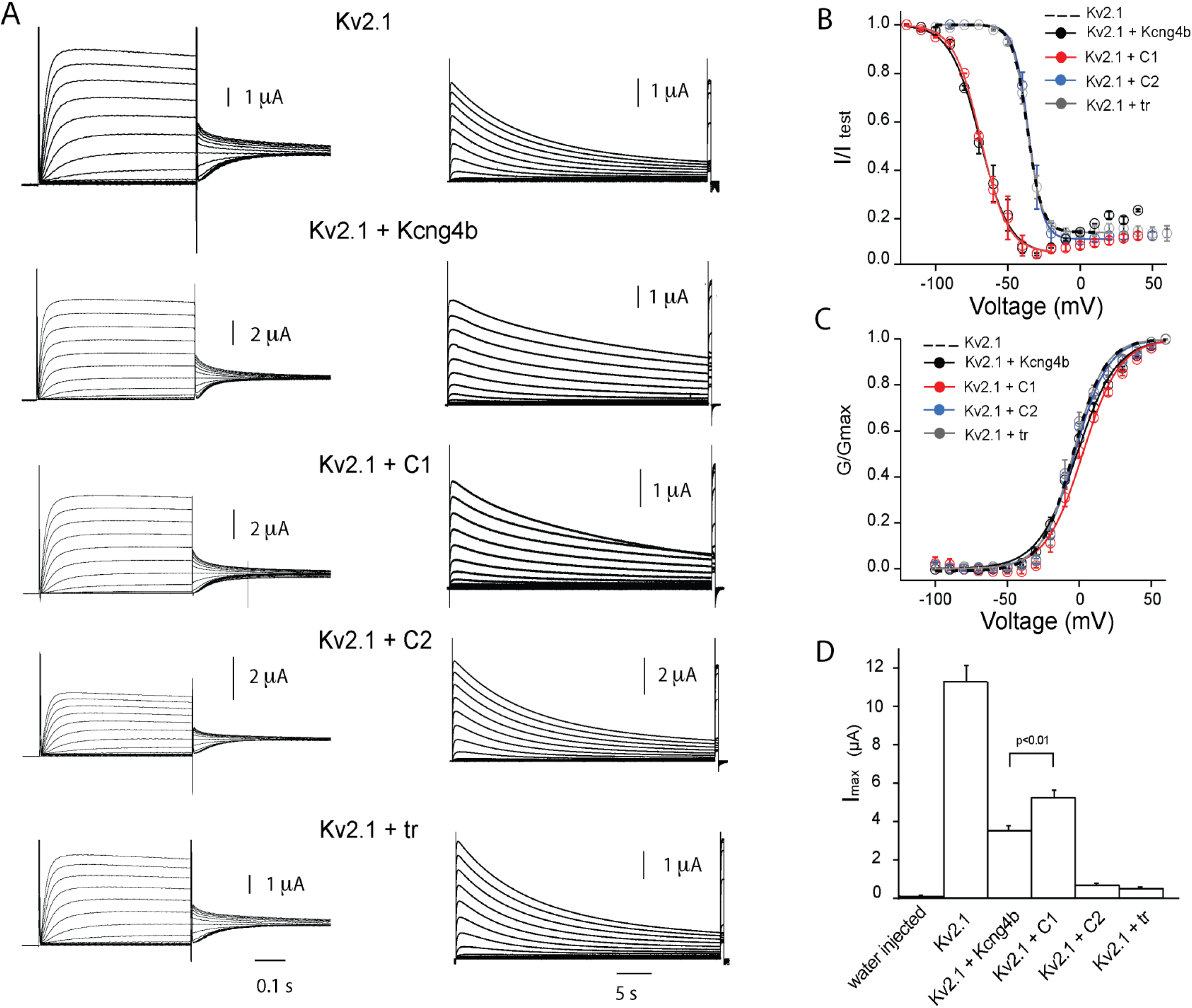
Biophysical properties of homomeric Kv2.1 (KCNB1) and heteromeric Kv2.1/Kv6.4 (Kcng4b) channels studied in *X. laevis* oocytes. A - Whole-oocyte current traces of Kv channels elicited by step-depolarization of the oocyte membrane from -100 mV to +60 mV with 10 mV increments (left) for estimation of steady-state activation. Current traces for estimation of the steady-state inactivation of channels (right). Current was induced by step-depolarization of membrane from -100 mV with 10 mV increments followed by short (300 ms) test pulse to +60 mV. The holding potential of oocytes expressing Kv2.1+kcng4b-wt and Kv2.1+kcng4b-C1 was - 120 mV in inactivation protocol. B - Steady-state inactivation curves of channels. C - Conduction-voltage (GV) relationship of channels shown in A. D - Current density of channels measured as peak current +60 mV. Error bars in the graphs represent ± SEM. Solid curves fit the data to Boltzmann function. P>0.01 – Student’s unpaired T-test was used to compare the groups. n≥ 7 for all experiments. Abbreviations: C1 - kcng4b-C1; C2 - kcng4b-C2; tr - N-terminal truncated Kcng4b (Shen et al., 2016)

In contrast, a dramatic reduction of current density was observed upon co-expression of KCNB1 with both Kcng4b-C2 and Kcng4b-trunc (Fig. 4D). Furthermore, the biophysical properties of channels formed by co-injection of KCNB1 with these Kcng4b variants were almost identical to those of KCNB1 homomers (Fig. 4; Table 1), indicating that the measured currents were generated by KCNB1 homomers. Since the current amplitudes of oocytes co-injected with KCNB1/Kcng4b-C2 and KCNB1/Kcng4b-trunc combinations are approximately 1/16 of the current mediated by homomeric channel (Fig. 4D), we consider it likely that these Kcng4b mutant forms exert a dominant-negative effect on Kv2.1 activity in the *Xenopus* oocyte system. A similar dominant-negative effect, but with much lesser extent, was previously shown by co-expressing Kcng4b-trunc with KCNB1 in HEK 293 cells (Shen et al., 2016). This inconsistency likely reflects the difference between the *Xenopu*s oocyte and HEK cells expression systems. The former allows better control over the ratio of expressed proteins. Thus, the simplest explanation for the substantial current reduction in oocytes co-injected with the stabilized mRNA of KCNB1/Kcng4b-C2 and KCNB1/Kcng4b-trunc is that heterotetramers containing Kcng4b mutant variants become much less functional or completely non-functional. Since the Kcng4b-trunc polypeptide lacked the protein domains forming ion permeation pore, but retained the tetramerization domain (Fig. 3), the observed dominant-negative effect can be explained by the formation of ion-conducting defective heterotetramers.

The nature of the dominant-negative effect of Kcng4b-C2 on KCNB1 could be more complex due to much more significant structural changes in the Kcng4b-C2 protein. The non-conducting heterotetramers could be formed or the KCNB1 inhibition could be increased due to the appearance of S7 TM domain. The resulting effect of the two mutants on Kv2.1 activity detected in *Xenopus* oocytes could be as follows. Kcng4b-C1 likely acts as a weakened Kv2.1 modulator with partial LOF, resulting in the Kv channel GOF, whereas Kcng4b-C2 strongly inhibits Kv2.1, which results in the Kv channel LOF.

### *kcng4b* mutants develop abnormal ears

The *kcng4b-C1^-/-^* embryos exhibited cell delamination in the third ventricle and mild hydrocephaly at 28 hpf (Suppl. Fig. 3A-D) along with abnormality of the heart, peripheral vasculature (Suppl. Fig 3, E, F) and ear. Thus, the phenotype of *kcng4b-C1* mutant is similar to that of the *kcng4b-trunc* mutant (Shen et al., 2016). The ear development was affected in the *kcng4b-trunc* mutant, where the ear size and number of otoliths increased (Suppl. Fig. 1). In contrast, in the *kcnb1* mutant the ear size and number of otoliths were reduced (Jedrychowska et al., 2021). In the *kcng4b-C1^-/-^* mutants, the ear was enlarged: at 28 hpf [7.3 ± 5.7 %, p = 0.01 (20 embryos, left + right ear)]; at 72 hpf [19.8 ± 7.9 %, p = 0.0001 (20 embryos, left + right ear)].

The additional third otolith was detected in abnormally high proportion during the first two days of development (Fig. 5C) like that in *kcng4b-trunc* (Suppl. Fig. 1), but later it disappeared, likely due to a fusion with the saccular otolith (Fig. 5F).

**Fig. 5.**
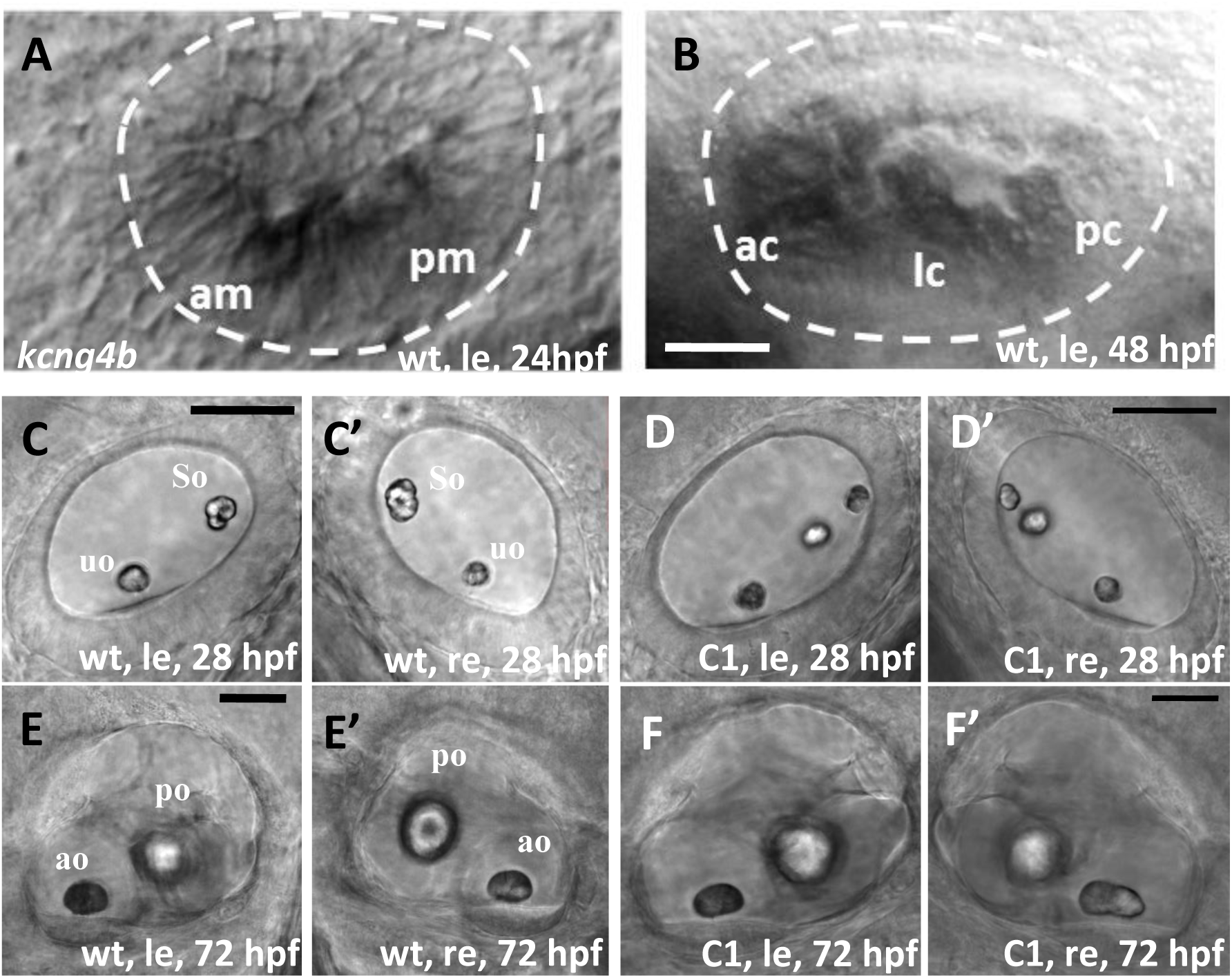
Kcng4b plays a role in ear development. All images are lateral views (anterior towards the left). A, B – whole mount in situ hybridization, *kcng4b*, left ear, A - 24 h; B - 48 hpf. C- F - DIC microscopy. C-E – ears at 28 hpf; and D, F - 72 hpf, le and re – left and right ears, uo and so – utricular and saccular otoliths, correspondingly; A, B, C, E – wildtype, D, F – *kcng4b-C1* mutant. Abbreviations: C1 - *kcng4b-C1*; hpf - hours post fertilization; le - left-hand side ear; re - right-hand side ear; wt - wildtype. Scale bar = 50 µm.

Unlike *kcng4b-C1^-/-^* mutants, *kcng4b-C2^-/-^* mutants develop relatively normally (Fig. 6A- D) except for the ears, where the opposite “no otolith” phenotype was observed for 8 days (Fig. 6E-H).

**Fig. 6.**
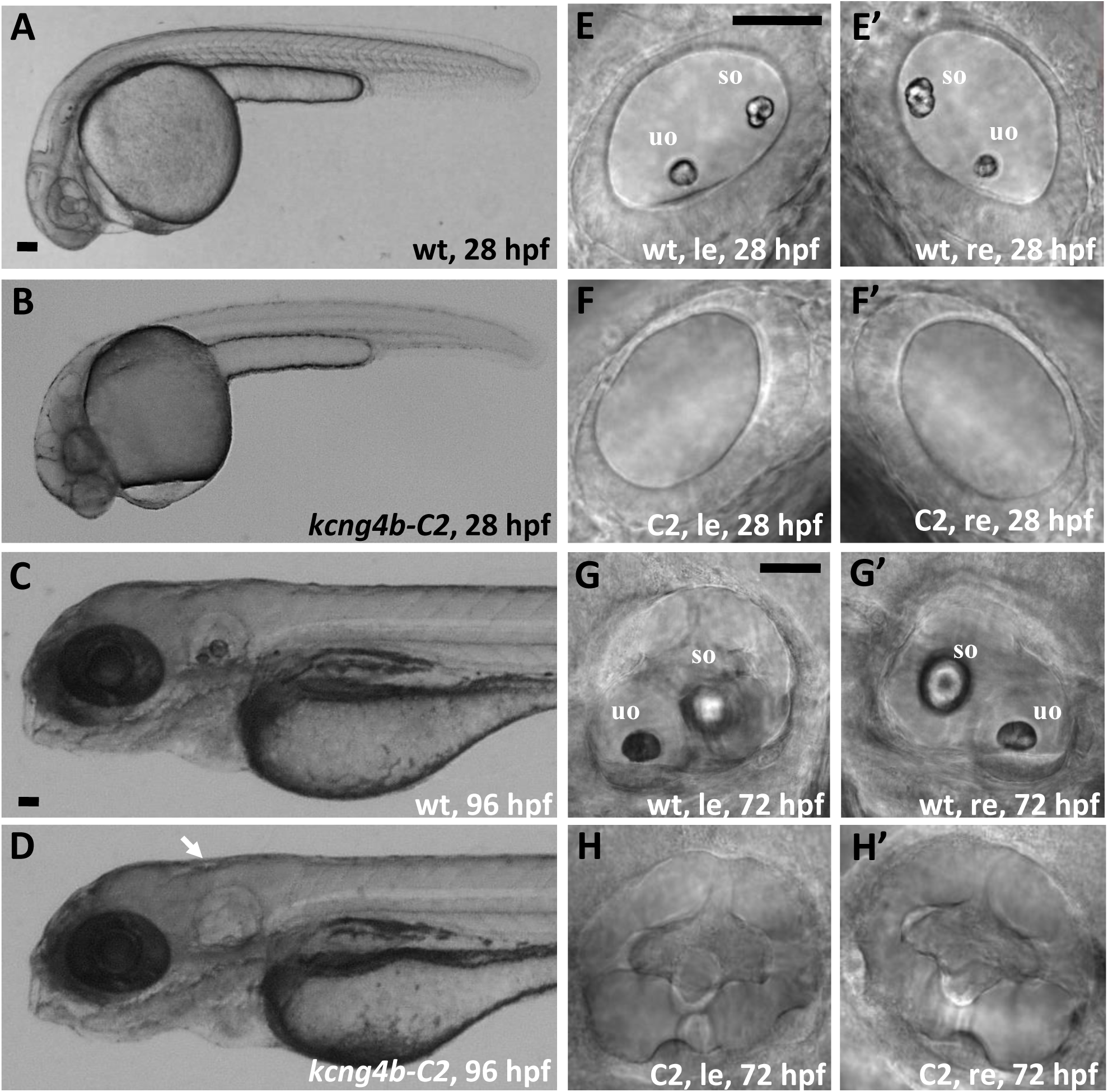
*kcng4b-C2* mutation causes an absence of otoliths. A, B – 28 hpf, arrow - ear; C, D – 96 hpf; E-H – ears at 28 (E, F) and 72 hpf (G, H), le and re – left and right ears, uo and so – utricular and saccular otoliths, correspondingly. All images are in lateral view. A, C, E, G – wild type, B, D, F, H – *kcng4b-C2*. Abbreviations: C2 *- kcng4b-C2*; hpf - hours post fertilization; le - left-hand side ear; re - right-hand side ear; wt - wildtype. Scale bar = 50 µm.

### *kcng4b* mutations affect kinocilia of sensory cells

The abnormal development of the ears may include defects of patterning and/or differentiation of sensory cells (Jedrychowska et al., 2021). In *Tg(Brn3c:GAP43-GFP)^s356t^*embryos, EGFP is expressed in the ciliated sensory cells of the ear and lateral line (Xiao et al., 2005). Analysis of *kcng4b-C1^-/-^/Tg(Brn3c:GAP43-GFP)^s356t^*and *kcng4b-C2^-/-^*/*Tg(Brn3c:GAP43- GFP)^s356t^* composite embryos and larvae (28 hpf and 72 hpf) did not reveal any obvious defect in the number or morphology of ear sensory cells (Suppl. Fig. 4, 5). However, the kinocilia of the sensory cells were affected. In 28 hpf *kcng4b-C1^-/-^* mutants, kinocilia morphology was mildly affected, and their number decreased (Fig. 7B, D), whereas in the 28 hpf *kcng4b-C2^-/-^* mutants, the kinocilia size and number were severely reduced (Fig. 7C, D).

**Fig. 7.**
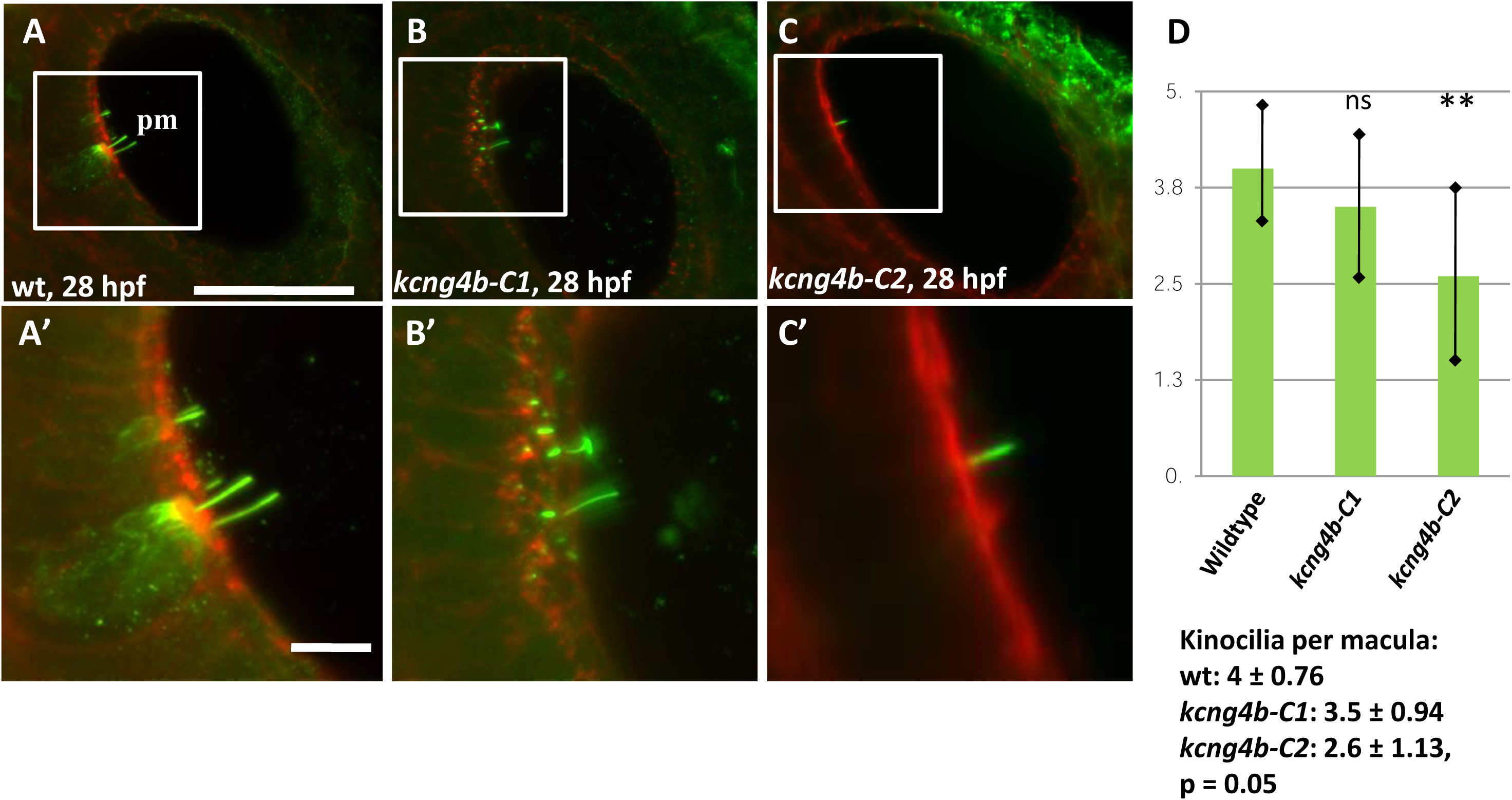
The development of inner ear sensory cells is affected in *kcng4b-C1* and *kcng4b-C2* mutants. Immunohistochemical staining using the anti-acetylated tubulin antibody (green) / phalloidin (red), 28 hpf, right-hand side ear. A – wild type, B - kcng4b-C1, C - kcng4b-C2, D – number of kinocilia per macula.

### Kcng4b mutations affect expression of multiple genes that regulate ear development

We have previously shown that the Kcnb1 LOF causes changes in gene expression (Jedrychowska et al., 2021). Here, we asked whether it is also true for mutations affecting the modulatory “silent” subunit of Kv2.1, Kcng4b. The two LOF mutants *kcng4b-trunc* and *kcng4b-C1* show rather similar phenotypes: enlarged ear and increased number of otoliths. In contrast, the *kcng4b-C2* mutants show the small ears and “no otoliths” phenotype. This phenomenon could be faithfully reproduced using the morpholino-or CRISPR-mediated knockdown (Shen et al., 2016). Aiming to address a cause for developmental changes in these mutants, we started with the comparative RNAseq analysis of transcriptomes of the Kcng4b-deficient embryos (crispants and morphants) by RNAseq (Suppl. Fig. 6). This analysis revealed changes in transcription of several genes known to play a role during development of the ear and otoliths.

These results were used as a guide to select the candidate genes for qRT-PCR analysis of the *kcng4b-C1* and *-C2* mutants at 24, 48 and 72 hpf (Suppl. Table 1). We focused on genes expressed in the mechanosensory hair cells, sensory patches, and otic vesicles. First, genes whose expression was affected in the same way at all three developmental stages (marked in bold black font in the Suppl. Table 2-4). These included *atp1b2b, atp2b1a, cldn7b, dachb, myo6b, sox9a, i.e.,* the genes expressed in the mechanosensory hair cells and otic vesicle. Second, the genes, whose expression varied from stage to stage, suggesting feedback autoregulation (marked in bold magenta in the Suppl. Table 2-4) - *igfbp3, kcng4b, mif, otofa, otogl, otop1, prom1a, sparc, wwc1.* Most of these genes were associated with otoliths formation. Third, *cldn7b, fgf3, fgf8a, gas8, gsdmeb, hapln1a, jag1* were expressed in two mutants in opposite ways during at least one developmental stage, which seems to be a common theme for changes in expression of all genes.

Previously, it was shown that in Kcnb1 mutants, *otop1* expression increased in parallel with reduction in otolith number (Jedrychowska et al., 2021). According to the qRT-PCR results, *otop1* expression was significantly affected in the Kcng4b mutants. The whole mount *in situ* hybridisation analysis of *otop1* expression supported these results and showed that the level of *otop1* expression changed dramatically. It was decreased in the *kcng4b-C1* mutants (Fig. 8A, B) and increased in the *kcng4b-C2* mutant (Fig. 8A, C), like that in the Kcnb1 LOF mutant.

**Fig. 8.**
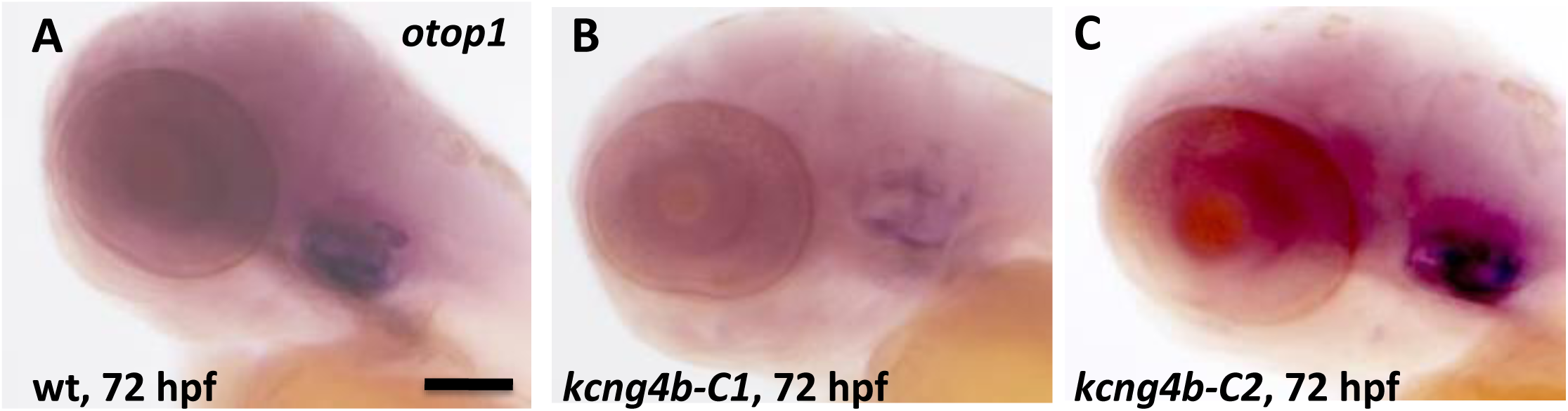
The expression of *otop1* was affected in *kcng4b-C1* and *kcng4b-C2* mutants as detected by whole mount in situ hybridization at 72 hpf. A – wildtype, B - *kcng4b-C1*, C - *kcng4b-C2*, Scale bar, 100 µm.

## Discussion

The plasma membrane (PM) potential (Vm) plays a role in cell survival, proliferation and regulation of transcription (Levin, 2014). Modulation of the electrically active KCNB1 by the electrically-silent KCNG4 controls channel activity (Bocksteins, 2016). The interaction of Kcnb1 and Kcng4b regulates cell survival, proliferation and development of zebrafish BVS, where Kcng4b LOF or Kcnb1 GOF increases and Kcng4b GOF or Kcnb1 LOF decreases Kv channel activity (Shen et al., 2016). The data presented illustrate a significant difference in inner ear development caused by mutations affecting the Kcng4b C-terminal.

The difference may be due to the structural difference between the putative mutant proteins. Kcng4b-C1 represents a milder version of the mutation causing Kcng4b LOF, which according to electrophysiological data (Table 1) is slightly less efficient in modulating the channel compared to the wildtype Kcng4b (Table 1), likely resulting in the mild channel GOF. Surprisingly, such a small increase in channel activity triggers a rather distinct developmental effect (Fig. 5), in indication of importance of precise tuning of channel activity by the modulatory subunits. Moreover, the mild LOF effect *in vitro* is further enhanced by the progressive reduction of Kcng4b-C1 transcript *in vivo* (Suppl. Table 2-4). Thus, the developmental effect may represent the sum of the mild LOF and *kcng4b* transcript reduction.

In contrast, the ectopic TM domain Kcng4b-C2 causes a dominant negative effect on Kcnb1 *in vitro* (Table 1), which alone can strongly block Kv channel activity. This effect seems to persist even at reduced transcript level (Suppl. Table 2, 3). Furthermore, the level of this transcript seems to increase in larvae up to 72 hpf (Suppl. Table 4). Of interest is a difference in the functional effects of the two potentially inhibitory forms of Kcng4b - Kcng4b-trunc and Kcng4b-C2 (Table 1), which cause opposite developmental effects (Fig. 6; Suppl. Fig. 1). This is most likely because the Kcng4b-trunc is unstable resulting in the Kv channel GOF (Shen et al., 2016) unlike dominant-negative Kcng4b-C2, which causes Kv channel LOF.

The electrophysiology data (Fig. 4, Table 1) suggest that Kcng4b-C2 suppresses the current amplitude of Kv2.1 like Kcng4b-trunc. Thus, despite structural changes caused by the mutation in the C-terminal by the appearance of ectopic 7TM domain, at least some interaction may take place between Kcnb1 and Kcng4b-C2. The remaining current kinetics are almost identical to those of Kcnb1 homomers, suggesting that interaction with Kcng4b-C2 renders the channel non-functional. This contrasts with Kcng4b-C1, which has similar but functionally weaker characteristics compared to Kcng4b-wt.

These data suggested that a fraction of Kcnb1 homomers remains active in the *kcng4b-C2* mutant. It may explain the lack of phenotype in the brain (Fig. 5), where the activity of the Kcnb1 homomer fraction may be just sufficient to avoid cell delamination. In contrast, in the ear, which is known to have a much higher concentration of K ions, this activity may not be sufficient to maintain otoliths development.

The KCNB1 human GOF phenotypes have been associated with epileptic encephalopathy, whereas the KCNB1 LOF phenotypes appear to cause developmental delay and speech defects (Bar et al., 2020; Latypova et al., 2017; Marini et al., 2017; Veale et al., 2022). Previously, our study demonstrated a reduction in otic vesicle size and number of otoliths associated with hearing and balance defects in zebrafish *kcnb1* LOF mutants (Jedrychowska et al., 2021). In the absence of otoliths in Kcng4b-C2, the developing zebrafish is likely to lack proper balance, as it was shown for embryos deficient in otoliths/otoconia (Hughes et al., 2004; Jedrychowska et al., 2021; Lundberg et al., 2015; Söllner et al., 2004). This illustrates that the role of Kcnb1-Kcng4b axis in the inner ear development is conserved in evolution between teleosts and mammals.

Language development is closely linked to hearing (Anne et al., 2017). In some human heterozygotic patients carrying a KCNB1/Kv2.1 mutation, speech development is impaired (Latypova et al., 2017; Marini et al., 2017; Bar et al., 2020). The current study of the two *kcng4b* mutants shows that different mutations affecting the silent α-subunit of the Kv channel can cause both LOF and GOF of the subunit, like what has been shown for the human electrically active subunit KCNB1. Given the antagonistic interaction between the two types of subunits (Shen et al., 2016), the LOF and GOF mutations of Kcng4 may result in the GOF and LOF effects on the functional heterotetramers of Kv, respectively.

## Acknowledgements

The authors thank Dr. Tanya Whitfield (University Sheffield, UK), all members of the Laboratory of Neurodegeneration and Zebrafish Core Facility of the International Institute of Molecular and Cell Biology in Warsaw for support, reagents and creative discussion.

VK acknowledges support from the Opus grants of the National Science Centre (NCN), Poland (2020/39/B/NZ3/02729).

## Author Contributions

JJ – analyzed mutants by immunohistochemistry and high-resolution microscopy and made Figures;

VV - analyzed mutants by electrophysiological analysis in *X. laevis* oocytes, made Figures, wrote and approved results;

MW, AM, JC – performed computer modelling, made Figures; wrote and approved results of computer modelling;

RA, RJ - performed qRT-PCR.

HS - performed analysis of kcng4b-trunc mutant phenotype and prepared RNA for RNAseq;

HC – performed detailed bioinformatics analysis of kcng4b-trunc RNAseq, made corresponding Figures and approved these results;

JK – designed experiments and discussed experimental results;

VK - designed mutants, supervised developmental biology experiments, analyzed RNAseq, made Figures, wrote and approved a paper.

## Competing Interests

The authors declare that they have no competing interests.**Table 1. Electrophysiological properties of K_v_2.1 homomeric channels as well as channels resulting from co-injection of K_v_2.1 with different forms of kcng4b protein.**

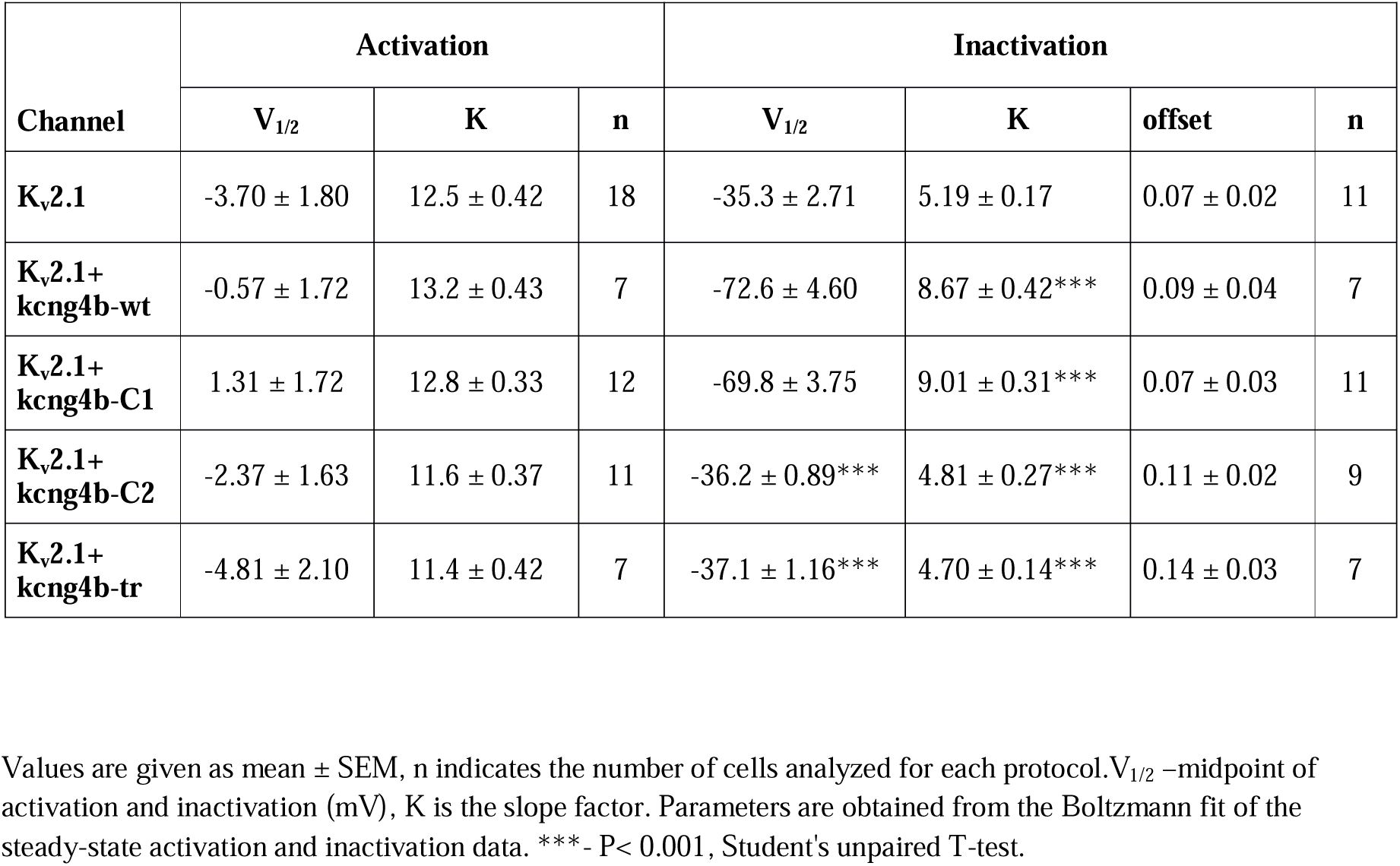

## Supplementary Materials

### Supplementary Figures Legends

**Supplementary Figure 1. The otic vesicle of *kcng4b-tr* mutants**. Note the changes in otoliths number and position (white arrow). The cell delamination in the third brain ventricle and death (white arrowhead) were described previously (Shen et al., 2016).

**Supplementary Figure 2. The 3D animation of structure of Kcnb1-Kcng4b-C2 mutant heterotetramer.** The additional TM domain of Kcng4b-C2 (S7) is in dark blue.

**Supplementary Figure 3. Phenotypically the kcng4b-C1 mutant is like the *kcng4b-tr* mutant** (Shen et al., 2016). All images are lateral views (anterior towards the left) except **D**, which is the front view. **A, B** – 28 hpf, arrowhead - IVth ventricle; **C, D** – cell delamination in the IIIrd ventricle (arrow); E, F – caudal vein deformation (marked by arrow), cv – caudal vein. **A, C, E,** – wild type, **B, D, F,** – kcng4b-C1. Abbreviations: hpf - hours post fertilization; le - left-hand side ear; re - right-hand side ear; wt - wildtype. Scale bar = 50 µm.

**Supplementary Figure 4. Patterning of sensory patches in the kcng4b-C1 mutant embryos of the Tg(Brn3c:GAP43-GFP)^s356t^) line is relatively normal. A** – left ear (le), 28 hpf; **B** – left ear, 72 hpf; **C** – lateral line (LL) tail neuromasts, 72 hpf. Ear vesicle’s boundary pointed by dotted line, am and pm – anterior and posterior maculae, ac, lc and pc – anterior, lateral and posterior cristae. **A, B, C** – wild type, **A’, B’, C’**, – *kcng4b-C1*. The scale bar is 50 micrometers.

**Supplementary Figure 5. Patterning of sensory patches in the *kcng4b-C2* mutant embryos of the *Tg(Brn3c:GAP43-GFP)^s356t^)* line is relatively normal. A** – left ear (le), 28 hpf; **B, C** – left ear, 72 hpf; **D –** lateral line (LL) tail neuromasts, 72 hpf. Ear vesicle’s boundary pointed by dotted line, am and pm – anterior and posterior maculae, ac, lc and pc – anterior, lateral and posterior cristae. **A, B, C, D –** wild type controls, **A’, B’, C’, D’ –** *kcng4b-C2*. Scale bar is 50 µm.

**Supplementary Fig. 6. The Kcng4b loss-of-function in zebrafish crispants and morphants at 24 hpf changes the expression of genes associated with ear and otolith development.**

