## Supplemental table 1 for "Mutant analysis of Kcng4b reveals how the different functional states of the voltage-gated potassium channel regulate ear development"

| N | **Genes** | **qRT-PCR primers, 5' - > 3'** |
| --- | --- | --- |
| 1 | *atp1b2b* | GCGGCGGTTTGAAGAACTTT  TCAGTCCACTCATCTTTCCCA |
| 2 | *atp2b1a* | CAGATCCTCTGGTTTAGAGG  TCCGAGTCCTCTATCCGG |
| 3 | *cldn7b* | GCCTTTGGATGTCGTGCGCT  GCCACACCCAGTCCAGCTACTG |
| 4 | *dachb* | CAGCGAGAGAATAAGGAAA  TGGAATGTTGTGAGGATG |
| 5 | *fgf3* | CCCAAGGGCGCCTTGTGCCAGGG  CCGTGTTTTAAAGCCCCTCCTGG |
| 6 | *fgf8a* | GCCGTAGACTAATCCGGACC  TTGTTGGCCAGAACTTGCAC |
| 7 | *gas8* | ACGAGTTTGCAAGGCCCATAA  TGGCCAACACTCTGTCCATT |
| 8 | *gsdmeb* | TATGAACTGTGTCTGCTG  ATCTGCTGGAATAACGAG |
| 9 | *hapln1a* | GGTGTGAGTTCATCGATGGG  CAAACGAGGGGAGTATGGGA |
| 10 | *igfbp3* | ATCTCACGGACACGACAAGC  GATGGACCGTCTCAGCTTCC |
| 11 | *jag1b* | GTGTGGATGGTGAAAACTGGT  GGGAGAAGACTGACACTCATTTATATT |
| 12 | *kcng4b* | ATTGCAGTGACAGTTATCAGC  AGCGCAACAAAAACTCCATGG |
| 13 | *mif* | GCGCAGAATAAACAATACTCC  TCCAAAGGTGCTGTTGTTCC |
| 14 | *myo6b* | TTTGAATGAGGCCACTCTCC  CAAAGAGCGTCCCTGATAGC |
| 15 | *otofa* | CACTGAAGTACACATGGAGC  GGTGACTTCAAAGCTGATGG |
| 16 | *otogl* | CAAAATGACCGAAGCGCACA  GCCCAGCTGTTCCCAAATTC |
| 17 | *otop1* | ATCTCTCAGGTACGGTACG  CGTCTCCTGTGGTTGTGC |
| 18 | *prom1a* | CGCTGCTGTGATAACTGTGG  CTCAGGTTCTGATTGGCTGC |
| 19 | *sox9a* | AGCACATCAGCTACGGTTCCTTCA  TCTGACCTCCAGCATGGGTGTAAT |
| 20 | *sparc* | TTGAGGTCGTGGAGGATGTT  GTATTTGCAGGGTCCGATGT |
| 21 | *stat3* | TGCCACCAACATCCTAGTGT  GCTTGTTTGCACTTTTGACTGA |
| 22 | *tecta* | CCTGGTATTGAGGGAGAGGAC  CGATCCACACCGTGTTCTTG |
| 23 | *tmie* | GGTGCAAAAGAAGGTAAAGAAGAGG  GCTTCTTTCTTTGGTGCTCCT |
| 24 | *ttc39c* | GGGAGAATAGATGAAGCC  GGAAGGTGAGGAATGGTA |
| 25 | *wwc1* | CGTTCTTCACTGTCTTCC  TTGTTTGTTGTCCTGGTAG |
| House-keeper | *eef1a1l1* | AAAATCGGTGGTGCTGGCAA  GGAACGGTGTGATTGAGGGA |
