## Supplemental table 2 for "Mutant analysis of Kcng4b reveals how the different functional states of the voltage-gated potassium channel regulate ear development"

Table 2. Comparison of expression of selected genes in the 24 hpf *kcng4b* mutants.

| N | Genes | **Expression level ratio,**  **kcng4b-C1 / wt, SD** | **p value** |  | **Expression level ratio,**  **kcng4b-C2 / wt, SD** | **Comment** |
| --- | --- | --- | --- | --- | --- | --- |
| 1 | *atp1b2b* | 0.34± 0.27 | 0.01 |  | 2.41±0.05 | hair cells, semicircular canal |
| 2 | *atp2b1a* | 0.05±0.90 | 0.05 |  | 7.04±0.008 | hair cells |
| 3 | *cldn7b* | 1.12±0.17 | 0.05 |  | 2.35±0.009 | otic vesicle, etc |
| 4 | *dachb* | 0.57±0.64 | n. s |  | 4.62±0.36 | otic vesicle, etc |
| 5 | *fgf3* | 2.97±0.59 | 0.05 |  | 1.87±0.005 | otic vesicle, etc |
| 6 | *fgf8a* | 2.80±0.63 | 0.05 |  | 1.65±0.02 | otic vesicle, etc |
| 7 | *gas8* | 2.76±2.58 | n. s |  | 4.43±0.002 | otic vesicle, not specific |
| 8 | *gsdmeb* | 1.79±0.18 | 0.01 |  | 1.28±0.05 | otic vesicle, not specific |
| 9 | *hapln1a* | 1.31±0.47 | n. s |  | 6.01±0.1 | semicircular canals, etc |
| 10 | *igfbp3* | 0.26±2.20 | 0.05 |  | 1.61±0.02 | otic vesicle, macula |
| 11 | *jag1b* | 0.17±0.60 | 0.001 |  | 1.72±0.008 | otic vesicle, etc |
| 12 | *kcng4b* | 3.43±0.79 | 0.05 |  | 0.63±0.02 | ependyma |
| 13 | *mif* | 1.47±0.02 | 0.0001 |  | 0.37±0.04 | otic vesicle, etc |
| 14 | *myo6b* | 0.2±1.64 | 0.05 |  | 5.4±0.006 | hair cells |
| 15 | *otofa* | 0.19±0.27 | 0.05 |  | 1.71±0.07 | hair cells |
| 16 | *otogl* | 0.33±0.72 | 0.05 |  | 1.71±0.08 | hair cells, otolith |
| 17 | *otop1* | 0.15±0.04 | 0.0001 |  | 3.17 ± 0.1 | otic vesicle, etc |
| 18 | *prom1a* | 1.76±1.40 | 0.01 |  | 0.76±0.12 | GFP-tagged in ET33-mi2a |
| 19 | *sox9a* | 0.38±0.82 | 0.05 |  | 5.27±0.09 | otic vesicle, etc |
| 20 | *sparc* | 0.64±0.35 | 0.05 |  | 1.88±0.12 | otic vesicle, etc |
| 21 | *stat3* | 0.30±0.91 | 0.05 |  | 0.55±0.009 | otic vesicle, etc. |
| 22 | *tecta* | 0.46±0.92 | 0.05 |  | 0.65±0.07 | hair cells |
| 23 | *tmie* | 1.17±0.06 | 0.001 |  | 1.55±0.09 | otic vesicle, etc |
| 24 | *ttc39c* | 1.18±1.08 | n. s |  | 7.48±0.07 | brain |
| 25 | *wwc1* | 1.44± 0.25 | 0.05 |  | 0.69±0.06 | not known |
