## Supplemental table 3 for "Mutant analysis of Kcng4b reveals how the different functional states of the voltage-gated potassium channel regulate ear development"

Table 3. Comparison of expression of selected genes in the 48 hpf *kcng4b* mutants.

| N | **Genes** | **Expression level ratio,**  **kcng4b-C1/ wt, SD** | **p value** |  | **Expression level ratio,**  **kcng4b-C2 / wt, SD** | **Comment** |
| --- | --- | --- | --- | --- | --- | --- |
| 1 | *atp1b2b* | 0.70±0.089 | 0.05 |  | 3.31±0.03 | hair cells, semicircular canal |
| 2 | *atp2b1a* | 0.57±0.32 | 0.05 |  | 3.07±0.07 | hair cells |
| 3 | *cldn7b* | 0.28±1.76 | n. s |  | 2.85±0.008 | otic vesicle, etc |
| 4 | *dachb* | 0.55±1.98 | n. s |  | 1.58±0.03 | otic vesicle, etc |
| 5 | *fgf3* | 1.21±1.52 | n. s |  | 0.82±0.005 | otic vesicle, etc |
| 6 | *fgf8a* | 0.83±0.15 | n. s |  | 0.88±0.06 | otic vesicle, etc |
| 7 | *gas8* | 2.64±1.12 | 0.05 |  | 2.06±0.01 | otic vesicle, etc |
| 8 | *gsdmeb* | 0.53±0.02 | 0.05 |  | 9.16±0.1 | not specific |
| 9 | *hapln1a* | 0.45±1.42 | 0.05 |  | 1.90±0.03 | semicircular canals, etc |
| 10 | *igfbp3* | 0.19±0.74 | 0.01 |  | 0.27±0.16 | otic vesicle, macula |
| 11 | *jag1b* | 0.17±0.60 | 0.01 |  | 0.27±0.08 | otic vesicle, etc |
| 12 | *kcng4b* | 0.60±0.28 | 0.05 |  | 0.68±0.03 | ependyma |
| 13 | *mif* | 0.51±0.21 | 0.05 |  | 5.21±0.18 | otic vesicle, etc |
| 14 | *myo6b* | 0.2±0.26 | 0.001 |  | 2.16±0.02 | hair cells |
| 15 | *otofa* | 2.46±0.72 | 0.05 |  | 0.47±0.05 | hair cells |
| 16 | *otogl* | 3.75±1.62 | 0.01 |  | 0.34±0.05 | hair cells, otolith |
| 17 | *otop1* | 0.04±0.001 | 0.05 |  | 2.57 ± 0.25 | otic vesicle, etc |
| 18 | *prom1a* | 0.62±0.31 | n. s |  | 0.61±0.14 | GFP-tagged in ET33-mi2a |
| 19 | *sox9a* | 0.33±0.55 | 0.05 |  | 7.45±0.22 | otic vesicle, etc |
| 20 | *sparc* | 1.50±0.21 | 0.05 |  | 3.56±0.01 | otic vesicle, etc |
| 21 | *stat3* | 0.44±0.4 | 0.05 |  | 0.60±0.001 | otic vesicle, etc. |
| 22 | *tecta* | 0.38±0.58 | 0.05 |  | 0.29±0.09 | hair cells |
| 23 | *tmie* | 2.07±0.12 | 0.05 |  | 0.48±0.07 | otic vesicle, etc |
| 24 | *ttc39c* | 5.00±0.50 | 0.05 |  | 0.63±0.04 | brain |
| 25 | *wwc1* | 0.6±1.62 | n. s |  | 2.21±0.16 | otic vesicle, etc |
