## Supplemental table 4 for "Mutant analysis of Kcng4b reveals how the different functional states of the voltage-gated potassium channel regulate ear development"

Table 4. Comparison of expression of selected genes in the 72 hpf *kcng4b* mutants.

| N | Genes | **Expression level ratio,**  **kcng4b-C1/ wt, SD** | **p value** |  | **Expression level ratio,**  **kcng4b-C2 / wt, SD** | **Comment** |
| --- | --- | --- | --- | --- | --- | --- |
| 1 | *atp1b2b* | 0.13± 1.54 | 0.01 |  | 9.57±0.003 | hair cells, semicircular canal |
| 2 | *atp2b1a* | 0.15±0.01 | 0.01 |  | 2.11±0.01 | hair cells |
| 3 | *cldn7b* | 0.75±0.48 | 0.05 |  | 1.54±0.001 | otic vesicle, etc |
| 4 | *dachb* | 0.07±0.83 | 0.01 |  | 1.62±0.01 | otic vesicle, etc |
| 5 | *fgf3* | 3.15±0.49 | 0.05 |  | 0.46±0.01 | otic vesicle, etc |
| 6 | *fgf8a* | 1.75±0.18 | 0.01 |  | 0.8±0.04 | otic vesicle, etc |
| 7 | *gas8* | 0.62±0.11 | 0.05 |  | 1.90±.026 | otic vesicle, etc |
| 8 | *gsdmeb* | 1.42±0.25 | 0.05 |  | 0.02±0.04 | not specific |
| 9 | *hapln1a* | 0.91±1.04 | n. s |  | 0.48±0.06 | semicircular canals, etc |
| 10 | *igfbp3* | 2.98±0.17 | 0.05 |  | 0.56±0.04 | otic vesicle, macula |
| 11 | *jag1b* | 0.25±0.4 | 0.01 |  | 0.54±0.07 | otic vesicle, etc |
| 12 | *kcng4b* | 0.28±0.22 | 0.01 |  | 7.05±0.01 | ependyma |
| 13 | *mif* | 0.76±0.17 | 0.01 |  | 0.01±0.02 | otic vesicle, etc |
| 14 | *myo6b* | 0.41±0.37 | 0.01 |  | 2.24±0.05 | hair cells |
| 15 | *otofa* | 0.37±0.43 | 0.01 |  | 1.11±0.03 | hair cells |
| 16 | *otogl* | 0.40±0.58 | 0.01 |  | 0.77±0.04 | hair cells, otolith |
| 17 | *otop1* | 0.06±0.006 | 0.001 |  | 1.44 ± 0.2 | otic vesicle, etc |
| 18 | *prom1a* | 0.66±0.76 | n. s |  | 2.19±0.01 | GFP-tagged in ET33-mi2a |
| 19 | *sox9a* | 0.73±0.80 | n. s |  | 1.50±0.1 | otic vesicle, etc |
| 20 | *sparc* | 0.34±1.51 | 0.05 |  | 1.02±0.04 | otic vesicle, etc |
| 21 | *stat3* | 0.47±0.30 | 0.05 |  | 0.34±0.04 | otic vesicle, etc. |
| 22 | *tecta* | 0.45±0.41 | 0.01 |  | 0.66±0.06 | hair cells |
| 23 | *tmie* | 0.76±0.20 | 0.01 |  | 1.21±0.07 | otic vesicle, etc |
| 24 | *ttc39c* | 0.26±1.54 | 0.05 |  | 2.52±0.04 | brain |
| 25 | *wwc1* | 0.76±1.32 | n. s |  | 0.59±0.22 | otic vesicle, etc |
