## Supplementary figures and images for "Mutant analysis of Kcng4b reveals how the different functional states of the voltage-gated potassium channel regulate ear development"

### Supplemental Figures

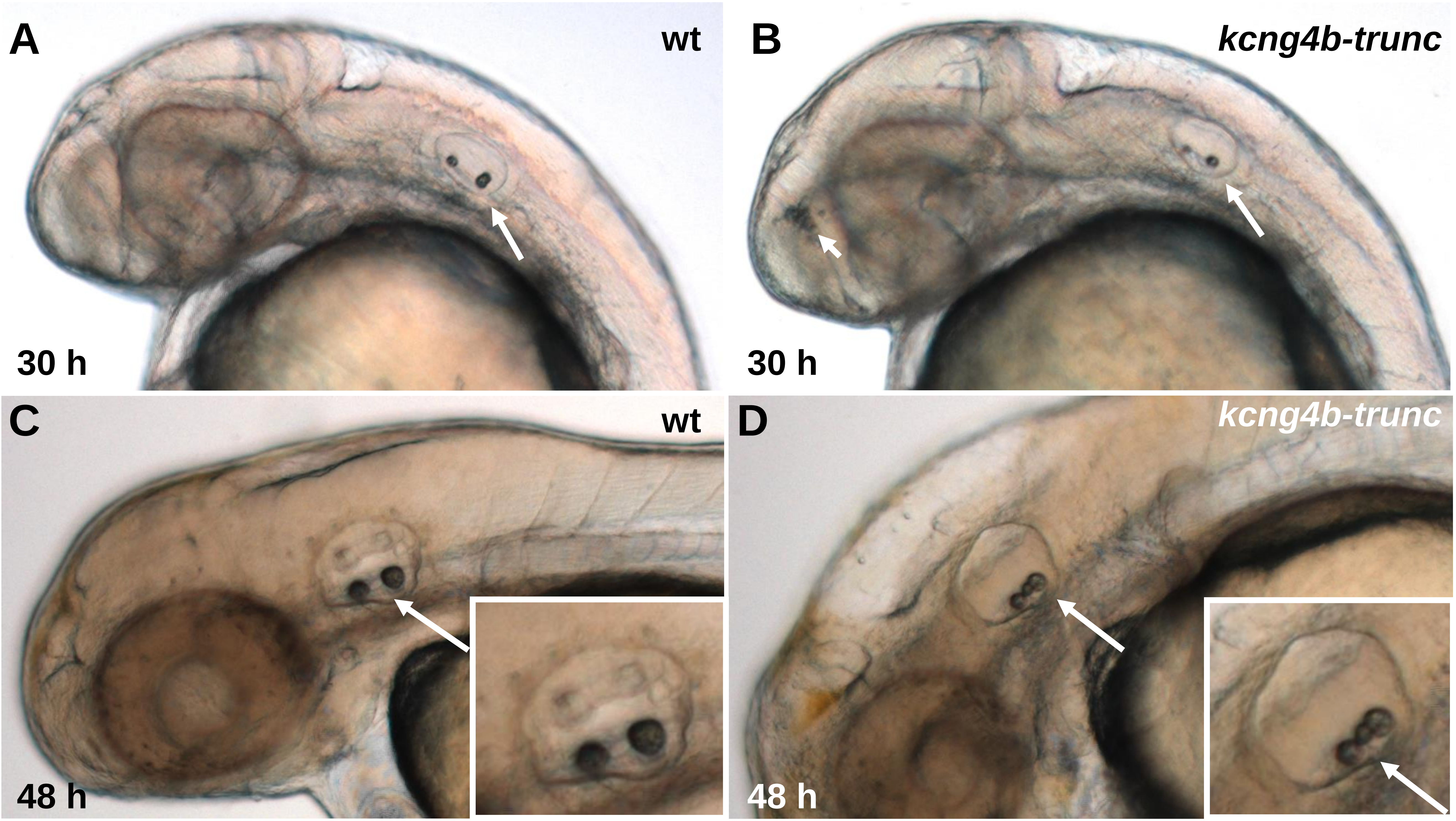

**Suppl. Fig. 2 (model animation will be provided during submission)**

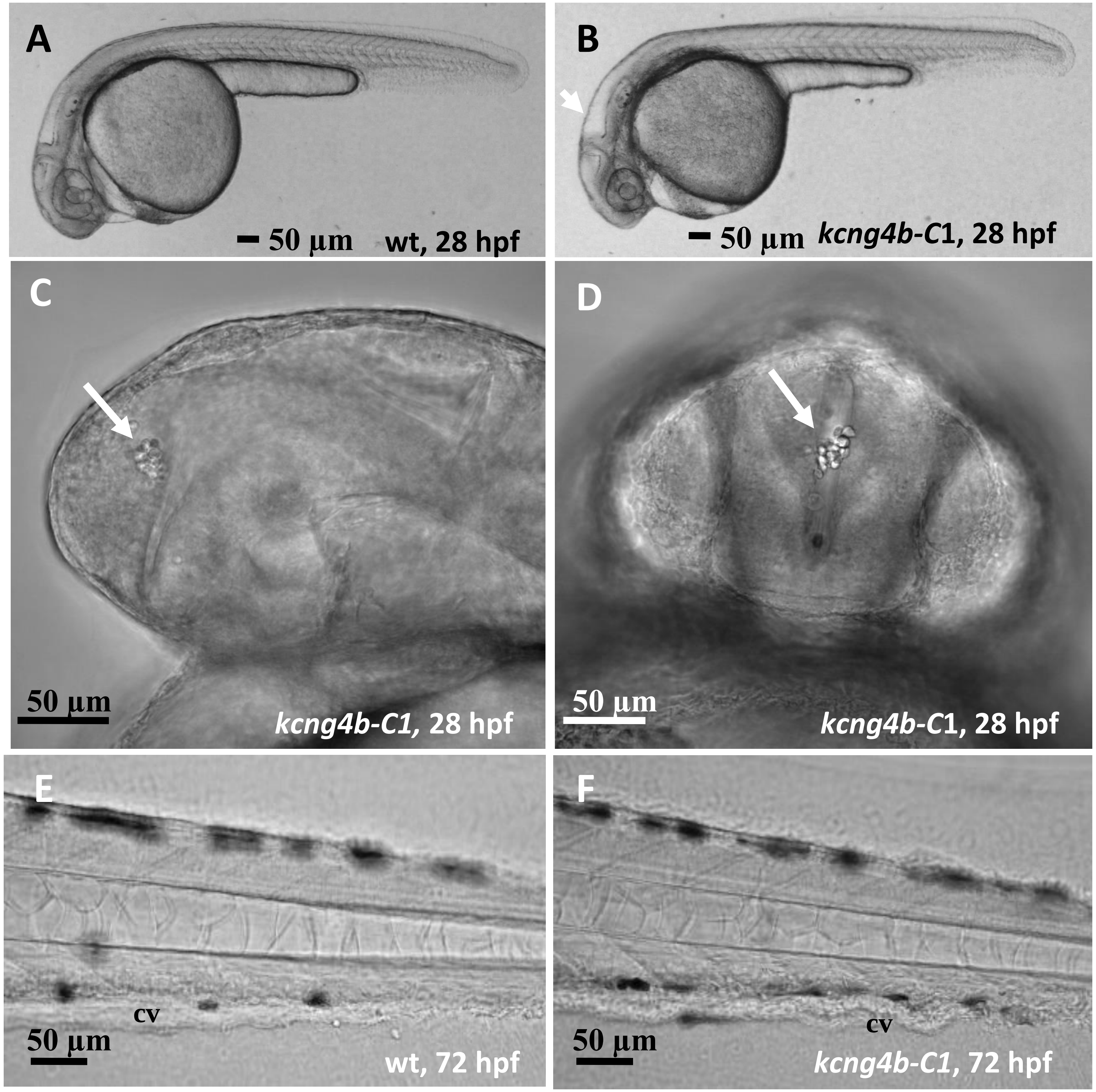

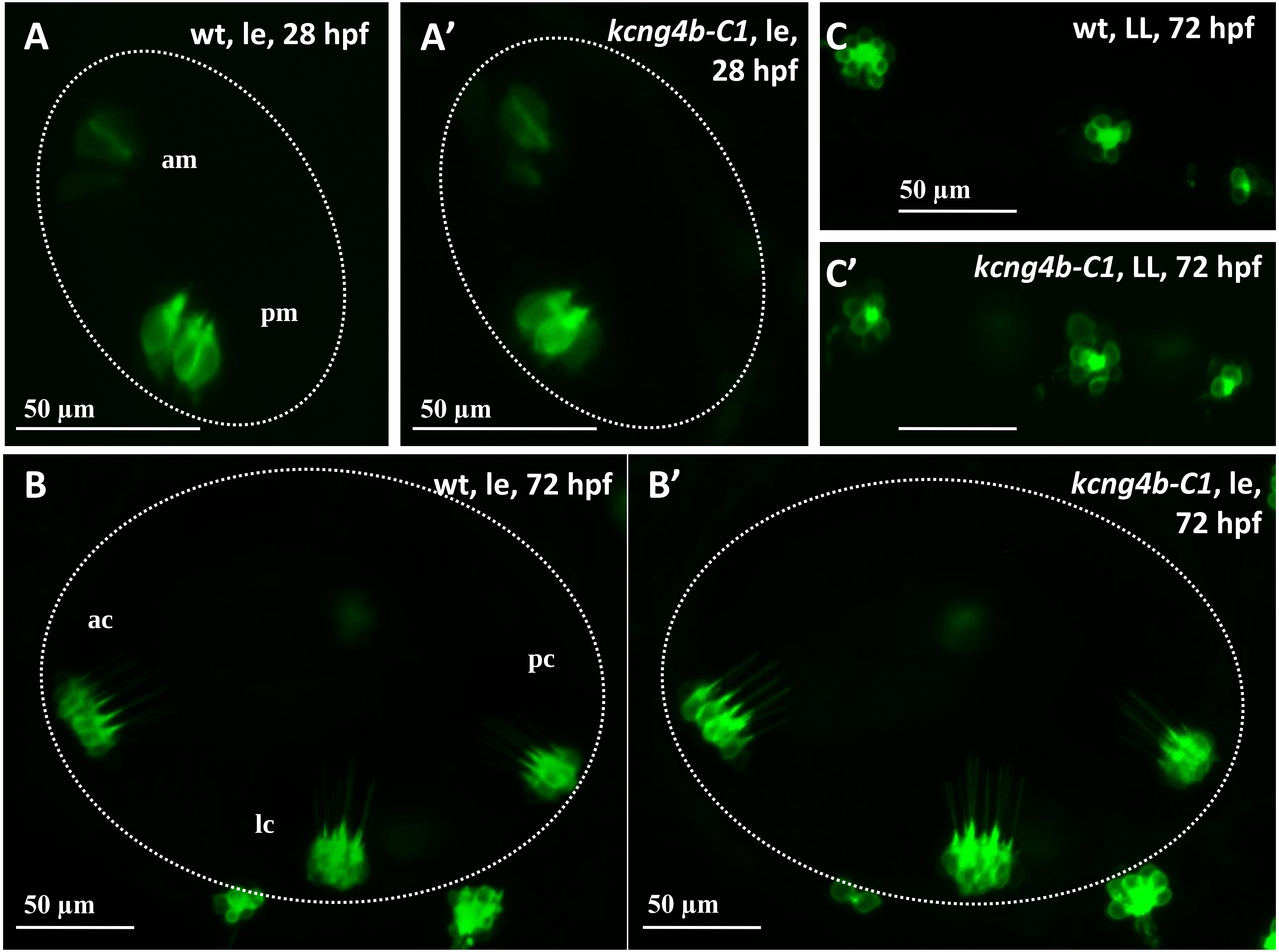

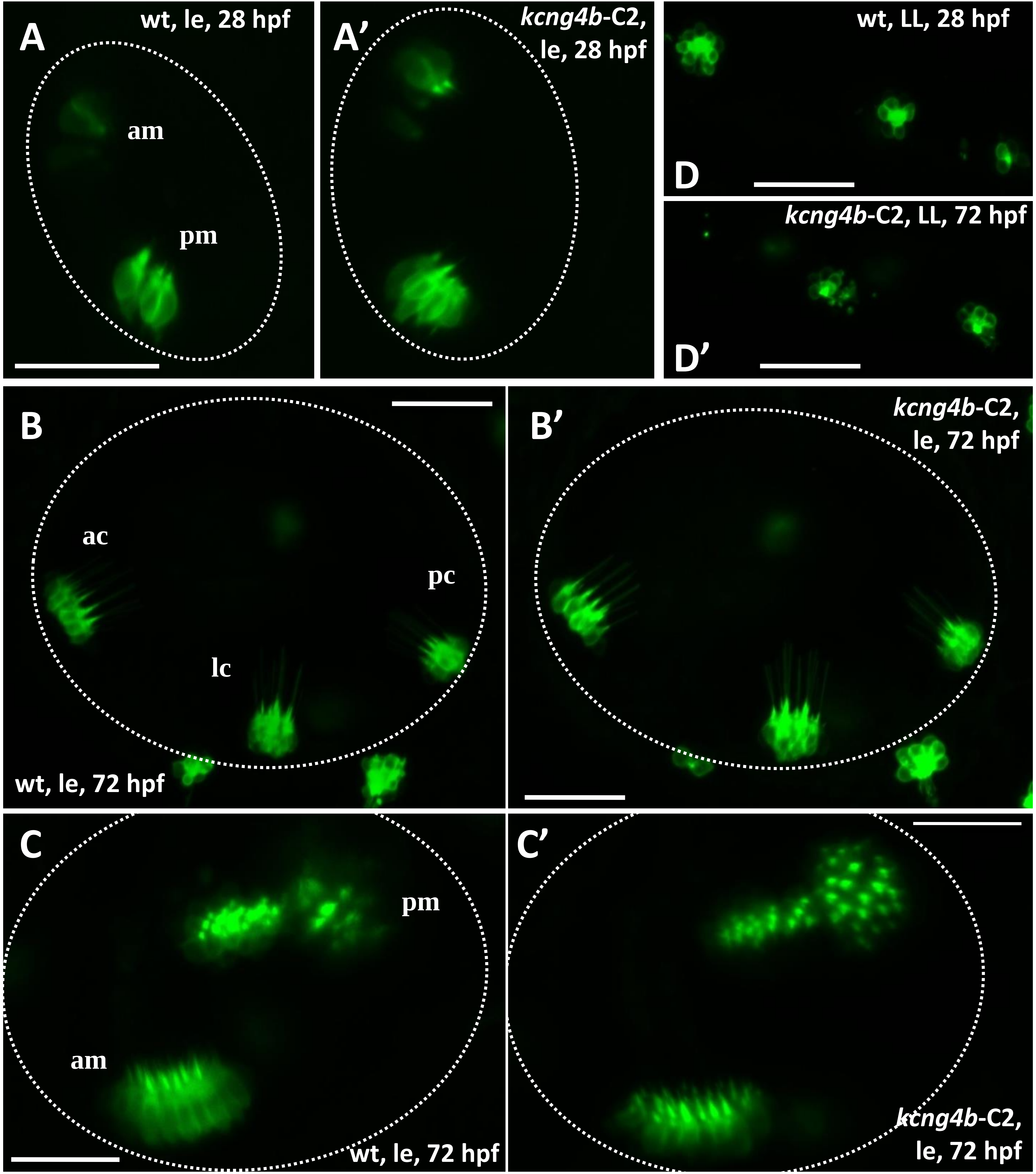

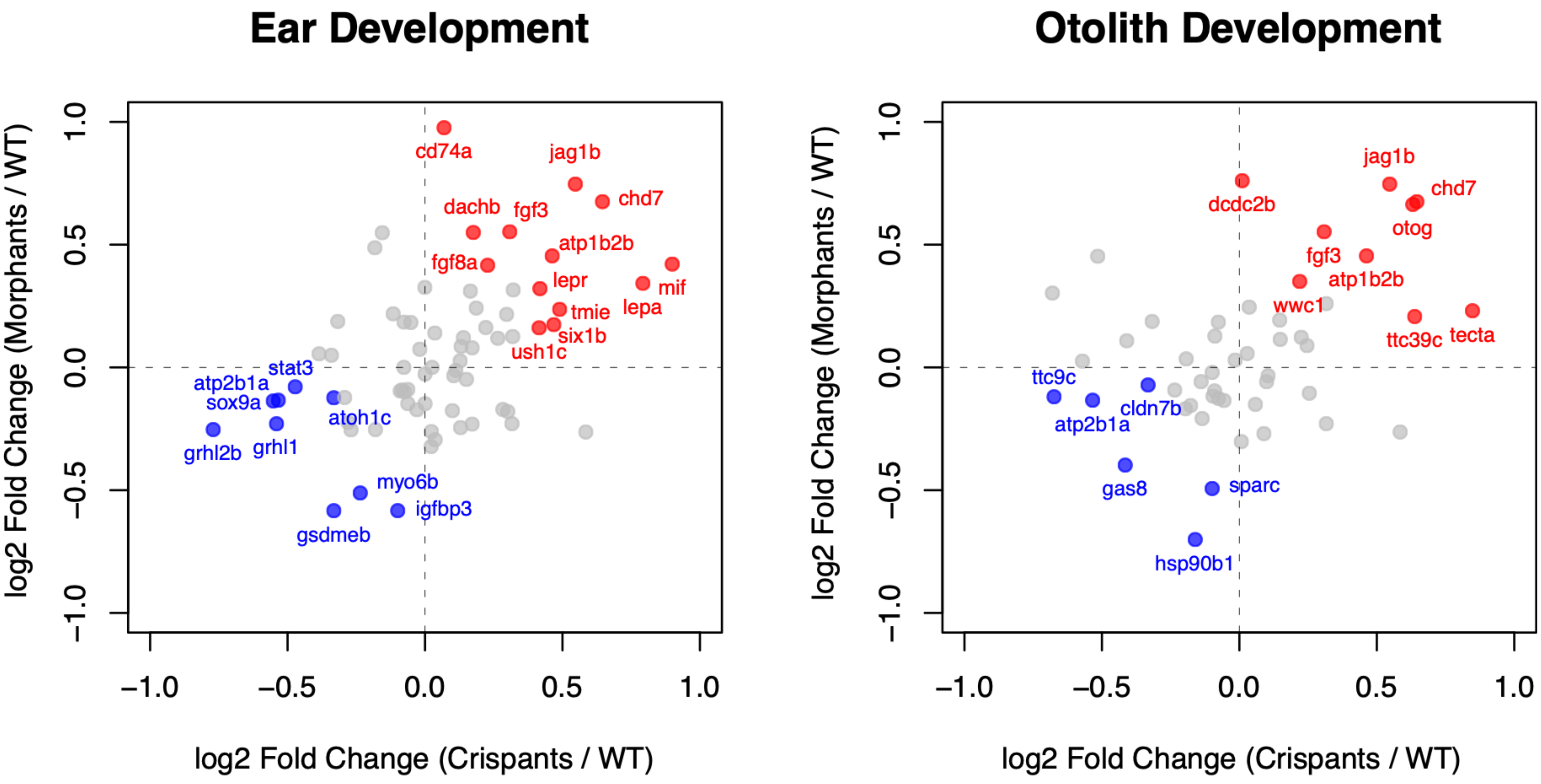
